# Cytokines mediate increased endothelial-leukocyte interaction and brain capillary plugging during CAR T cell neurotoxicity

**DOI:** 10.1101/2025.02.19.638920

**Authors:** Lina Park, Yu-Tung Tsai, Ruoqian Hu, Hyun-Kyoung Lim, Lila D. Faulhaber, Katelyn Burleigh, Mahashweta Bose, Eli M. Faulhaber, Isabella H. Draper, Stephen E.P. Smith, Andy Y. Shih, Alexandre Y. Hirayama, Cameron J. Turtle, Colleen E. Annesley, Rebecca A. Gardner, Heather H. Gustafson, Ying Zheng, Juliane Gust

## Abstract

CAR-T cells treat cancer, but also cause systemic cytokine release and immune effector cell associated neurotoxicity syndrome (ICANS). In an immunocompetent mouse model, we show by in vivo two-photon imaging that CD19-CAR T treatment causes brain capillary plugging by circulating CAR-T cells and other CD45+ leukocytes, as well as cortical hypoxia. This is accompanied by increased endothelial ICAM-1 and VCAM-1 expression in the brain capillary-venule transition zone, where most of the capillary stalls occur. In the mouse model, circulating CAR-T cells strongly upregulate integrin α4β1 affinity to VCAM-1, but not affinity of integrin αLβ2 to ICAM-1. Blockade of integrin α4 but not integrin αL improves locomotion behavior. In vitro, human brain microendothelial cells upregulate ICAM-1 more than VCAM-1 in response to TNF, IFN-γ, and IL-1β. In a 3D brain human microvessel model, treatment with TNF and IFN-γ is sufficient to induce adhesion of CAR T cells under flow conditions, which is blocked synergistically by antibodies against integrins α4 and αL. Finally, patients with the highest levels of TNF and IFN-γ also have the highest blood levels of soluble ICAM-1 and VCAM-1, which in turn correlate with ICANS. Integrin α4 but not αL increases in CAR-T cells after they are infused into patients. Combined data from patients, mouse models and in vitro microvessels indicate differential regulation of interactions of ICAM-1 and VCAM-1 with their respective leukocyte integrins. Overall, our study supports the hypothesis that cytokine-driven upregulation of endothelial-leukocyte adhesion is sufficient to induce acute, reversible neurotoxicity.

**One Sentence Summary:** During CAR T cell therapy, cytokine release induces white blood cell stalling in brain capillaries by upregulating ICAM-1/VCAM-1-integrin interactions.

## INTRODUCTION

Neurotoxicity has long been recognized as a characteristic adverse event following chimeric antigen receptor (CAR) T cell therapy for hematologic malignancies (*1*). The clinical syndrome has been termed immune effector cell associated neurotoxicity syndrome (ICANS) (*2*) based on the observation that different CAR products can cause a similar constellation of symptoms. The typical patient with ICANS will experience cytokine release syndrome (CRS) within the first week after CAR T cell infusion, followed by a decline in mental status a day or two later. Although these symptoms typically resolve without specific intervention, in some cases patients can rapidly progress to critical neurologic illness with seizures, coma, and even fatal cerebral edema (*1, 3*). Antecedent systemic cytokine release is a strong and consistently observed risk factor for ICANS (*4*). However, the cellular-molecular mechanisms of how this systemic inflammatory state causes brain dysfunction remain uncertain.

We have previously established an immunocompetent mouse model of ICANS (*5*). Wild-type non-tumor bearing mice received murine CD19-CAR T cells, which rapidly proliferate as they target normal CD19+ B cells everywhere in the body and lead to B cell depletion. This was accompanied by systemic cytokine release, behavioral changes, and brain microhemorrhages. Brain histology showed increased deposition of immunoglobulins, indicating loss of integrity of the blood-brain-barrier, loss of capillary pericytes, and formation of lumenless string capillaries (*5*). To understand how cerebral microvascular function is affected, we performed in vivo two-photon imaging through a thinned skull window. Surprisingly, we did not observe any leakage of intravascular tracer, but instead found extensive capillary plugging by circulating leukocytes that correlated temporally with the behavioral abnormalities, peaking 6 days after CAR T cell infusion (*5*). Similar brain capillary plugging has been observed in mouse models of acute ischemic stroke, seizure, and Alzheimer’s disease (*6–8*).

In humans, brain capillary size is 6-7μm (*9*) and leukocyte size is 6-8 μm (*10*). This tight fit forces the leukocytes to deform into an oblong shape to pass through brain capillaries. Even in normal conditions, leukocytes can briefly stall and cause decreased blood flow (*11, 12*). The increase in leukocyte stalling during inflammation could be explained by a decrease in overall blood flow, decreased capillary caliber, and/or increased adhesion between the leukocyte and the capillary endothelium (*13*).In our prior study, we found no change in capillary diameters from baseline, and no change in systemic blood flow after CAR T cell treatment (*5*). Leukocytes only obstructed capillaries less than 6μm in diameter. However, we also observed increased adhesion of leukocytes to the walls of vessels that were too large to become obstructed. This led us to our current hypothesis that systemic cytokine release is sufficient to induce upregulation of endothelial-leukocyte adhesion interactions to a degree that leads to dysfunctional adhesion to the brain endothelium, resulting in insufficient perfusion and cerebral hypoxia (*1, 13, 14*). An attractive feature of this hypothesis is that it also explains the predictable, monophasic and reversible course of ICANS, because the capillary plugging resolves as soon as the cytokine storm abates.

Using a combination of in vivo and in vitro models as well as patient biomarkers, we now demonstrate that systemic IL-1β, TNF, and IFN-γ increase brain endothelial ICAM-1 and VCAM-1 expression, and that CAR T cells increase the affinity of their integrin α4 to VCAM-1 after infusion into the body. Blocking the interaction of ICAM-1 and VCAM-1 with their respective integrins is sufficient to abolish CAR T cell adhesion to brain endothelium. In CAR T cell patients, these processes may be sufficient to induce clinically significant brain hypoperfusion due to capillary obstruction by stalled leukocytes.

## RESULTS

### CAR T cell capillary plugging is associated with focal brain hypoxia

We have previously validated an immunocompetent mouse model of ICANS, where treatment with murine CD19-CAR T cells in non-tumor bearing wild type mice induces systemic cytokine release with marked increases in IFN-γ and IL-6, and smaller rises in IL-1β and TNF (*5*). In a time course that mimics human ICANS, mice developed dose-dependent behavioral deficits. Using in vivo two photon imaging, we demonstrated plugging of >10% of brain capillaries by circulating leukocytes by day 6 after CAR T cell infusion (*5*).

We now used the same model to study the molecular underpinnings of brain capillary plugging. First, we determined whether the plugging leukocytes are primarily CAR T cells, or whether endogenous white blood cells also participate. We treated healthy wild-type mice cyclophosphamide lymphodepletion followed by 10 million tdTomato-coexpressing CD19-CAR T cells i.v. and imaged the cortical microvascular network by in vivo two-photon imaging through a chronic cranial window (Fig. 1A). The circulating leukocyte pool was labeled with intravenously injected fluorescently labeled anti-CD45 antibody. Of 145 nonflowing capillaries imaged between day 4-6 after CAR T cell infusion (N= 8 mice), 103 had visible plugs. When no obstruction was visible, this was typically because nonflowing capillaries were part of an obstructed network with the obstructing object outside of the field of view. 88% of the capillary plugs were CD45+ leukocytes. Both CAR T cells and non-CAR T leukocytes (Fig. 1, B and C) were observed to be plugging cortical capillaries. In some cases, the leukocyte plugs were associated with stalled red blood cells or microthrombi (Fig. 1C). However, the stalled red blood cells almost always appeared to be arrested behind a leukocyte plug that was obstructing blood flow, instead of being the primary cause of the obstruction. On occasion, CAR T cells could be observed crawling back and forth within the capillary, alternating with and against the direction of blood flow (Fig. 1D). This indicates active interaction between the T cell and endothelium rather than a passive mechanical process. Increased adhesion of CAR T cells and non-CAR T leukocytes was also observed in larger vessels, primarily dural and pial venules (Fig. 1E, Movie S1). We never observed active extravasation of any leukocytes into the brain parenchyma, whereas it was occasionally seen in venular hotspots in the dura (Fig. S1) Leukocyte adhesion or capillary plugging was only very rarely observed in healthy mice or controls treated with lymphodepletion only, or lymphodepletion plus mock-transduced T cells (*5*), and we never saw any crawling of cells against the direction of blood flow. We were not able to further specify the identity of plugging cells because other antibodies that we tested (directed against CD3, CD4, CD8, or Ly6G) led to rapid ablation of the target cells, precluding their use for stable in vivo labeling. CAR T cells were also present in the extravascular space, particularly in the meninges, but active extravasation events were observed only very rarely and only in the dural vasculature.

**Fig. 1.**
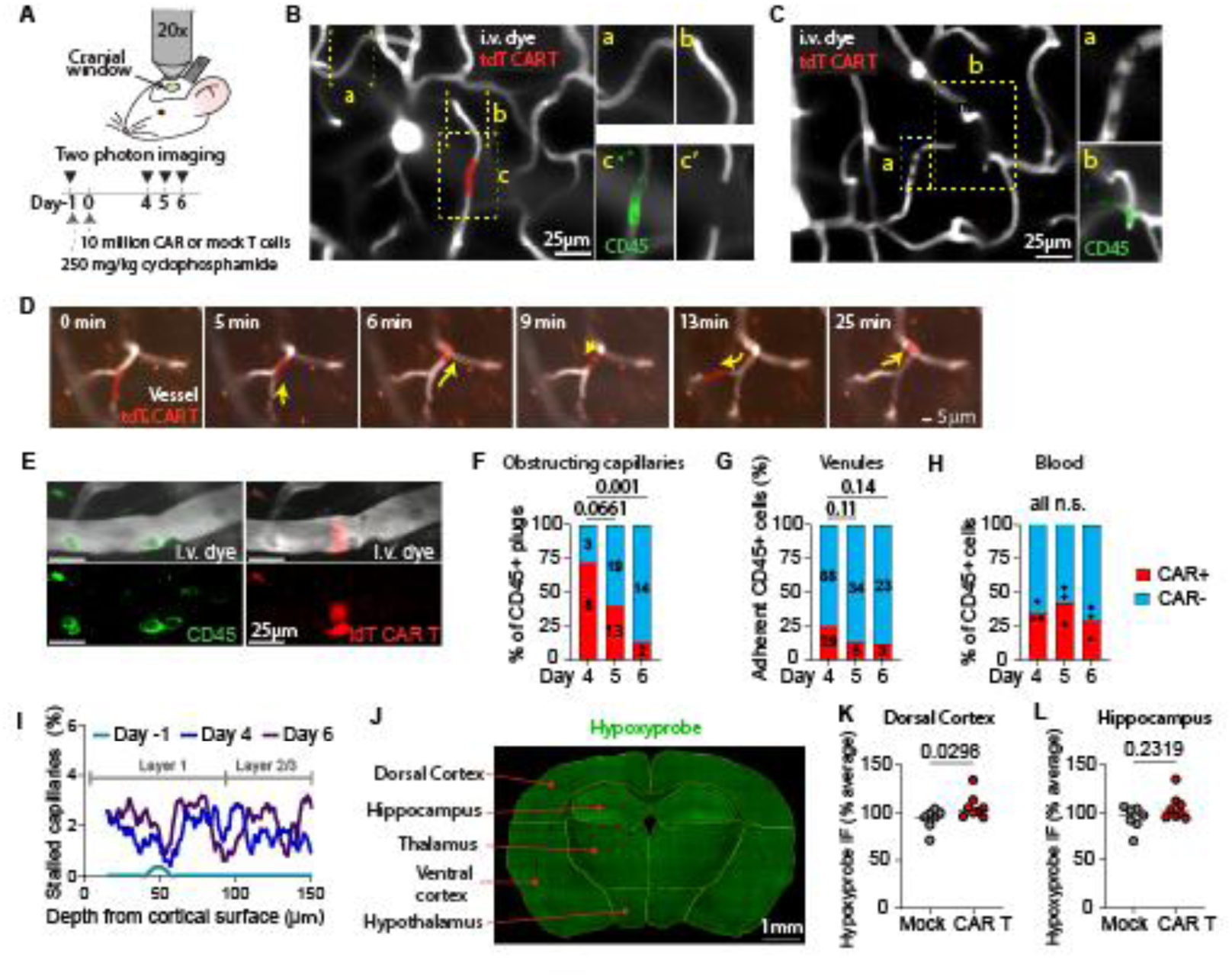
CAR T cells and non-CAR T leukocytes obstruct blood flow in brain capillaries during CD19-CAR T neurotoxicity. (**A**) Schematic of in vivo two photon imaging through a cranial window. Mice received 10 million CD19-directed CAR T cells cotransduced with tdTomato for in vivo imaging. (**B**) Representative example of a CAR T cell (expressing td-Tomato) obstructing a brain capillary, seen as top-down view through the cranial window. Insets: **a**, example of a flowing capillary, with striations created by transiting red blood cells. **b**, nonflowing capillary due to obstructing CAR T cell, note absence of striations. **c**, colabeling of CAR T cell with intravascularly delivered anti-CD45 antibody, and **c’** filling defect created by the CAR T cell. (**C**) example of a tdTomato-negative, non-CAR T leukocyte obstructing a capillary. Only a filling defect is seen in the red channel. Insets: **a**, discoid filling defects (∼5µm diameter) created by individual stalled red blood cells behind the leukocyte. **b**, CD45 labeling confirm that the obstructing object is a leukocyte. (**D**) Example of a stalled CAR T cell (red) moving back and forth within the capillary network over the span of 25 minutes. (**E**) Representative example of adherent leukocytes in a pial venule. The left and right images were acquired serially within 2 minutes of each other, because different excitation wavelengths were required for optimal acquisition of the CD45-Alexa488 and the tdTomato labels. Therefore, the cells have moved slightly between images. (**F**) Fraction of CAR T (tdTomato-positive) and CD45+, tdTomato-negative cells that were stalled and obstructing capillary flow, n=3-6 mice per timepoint. Numbers inside the bars show the number of cells evaluated, two-proportion z-test. (**G**) Fraction of CAR T and non-CAR T cells adherent to the luminal side of pial and dural venules, n=3-7 mice per timepoint. Numbers inside the bars show the number of cells evaluated, two-proportion z-test. (**H**) Fraction of CAR T cells and non-CAR T leukocytes in blood, each dot indicates one mouse, one-way ANOVA with Fisher’s LSD posttest. **(I)** Percentage of stalled capillaries by depth from cortical surface, values represent the average of 9-10 mice per timepoint. The lines show the running average (smoothened by 10 nearest neighbors on each side) of 1µm depth steps. The top 20µm is not shown to avoid edge artifact. (**J**) Coronal section of CAR T treated mouse brain stained for hypoxyprobe (green). The annotations show the regions that were manually segmented for analysis. (**K,L**) Scatter plots showing the mean brightness of hypoxyprobe immunofluorescence (IF) in the cortex and hippocampus. Values are normalized to the average for each cage (all cages had at least one CAR T and at least one mock control mouse). Each dot indicates one mouse, unpaired two-sided t test.

We next investigated whether the proportion of CAR T cells among plugging and adherent cells changes over time. On day 4 after CAR T cell injection, the first day we observed significant capillary plugging, CAR T cells made up 72% of all CD45+ capillary plugs (range, 60-100% for individual mice). This decreased to only 13% on day 6 (range, 0-18%) (Fig. 1F). This observation suggests that CAR T cells are first to become activated and increase their adherence to the brain endothelium, but then the immune activation spreads to other leukocyte subsets. To understand whether this shift in adherent populations is universal, we also examined the cell populations that adhere to the endothelium in larger venules of the brain border regions at the pial surface (Fig. 1G). Here, the proportion of CAR T cells also decreased over time, with CAR T cells representing 25% of all adherent leukocytes on day 4 (range, 11-50% for individual mice), and 12% on day 6 (range, 0-40%). The fraction of CAR T/total CD45+ cells in the blood differed from that of plugging cells, with 35% CAR T cells (range, 30-43%) on day 4 and 29% (range, 15-38%) on day 6 (Fig. 1H). This confirms that the fraction of plugging CAR T cells is not simply a reflection of their overall relative abundance. Together, these results support active regulation of leukocyte-brain endothelial adhesion during the peak of CAR T toxicity between day 4-6 after infusion.

Capillary plugging imaged through thinned-skull windows was very rare on day -1 (prior to CAR T cells). It substantially increased by day 4-6, and was relatively uniformly distributed across the first 150μm of cortical depth (Fig. 1I). To determine whether capillary plugging has detrimental effects on perfusion of the cortex or other regions that were inaccessible to our imaging approach, we performed hypoxyprobe labeling to detect hypoxic brain tissue. CAR T cell treated mice had a median 8% higher hypoxyprobe signal in the dorsal cortex compared to mice treated with mock transduced T cells (n=7-8 per group, P=0.0298) (Fig. 1, J-L). This corresponds to the brain region where we obtained our two-photon imaging data. There was no significant difference in hypoxyprobe labeling in other brain regions (hippocampus, thalamus, ventral cortex-striatum, and hypothalamus) between mock and CAR T treated mice.

### Upregulation of ICAM-1 and VCAM-1 in brain capillaries after CAR T cell treatment

To understand how leukocyte adhesion to the brain endothelium is regulated in the CAR T model, we first focused on intercellular adhesion molecule-1 (ICAM-1) and vascular cell adhesion molecule-1 (VCAM-1), the principal cell adhesion molecules that regulate neuroinflammation (*13*). We have previously shown that soluble ICAM-1 and VCAM-1 increase in the blood of immunocompetent mice after treatment with CD19-CAR T cells, and that ICAM-1 but not VCAM-1 increases in whole brain lysates of these mice (*5*). To determine if brain capillaries specifically upregulate these adhesion molecules, we first measured ICAM-1 and VCAM-1 expression in cortical capillaries by immunohistochemistry on brain sections. On day 6 after CAR T cell infusion, ICAM-1 was upregulated in cortical capillaries of CAR T cell treated mice compared to those receiving mock transduced T cells (n=4-6 per group, P=0.0095) (Fig. 2A,B). In contrast, VCAM-1 expression varied widely between individual mice, and was overall not increased over mock controls (P=0.9999) (Fig. 2C).

**Fig. 2.**
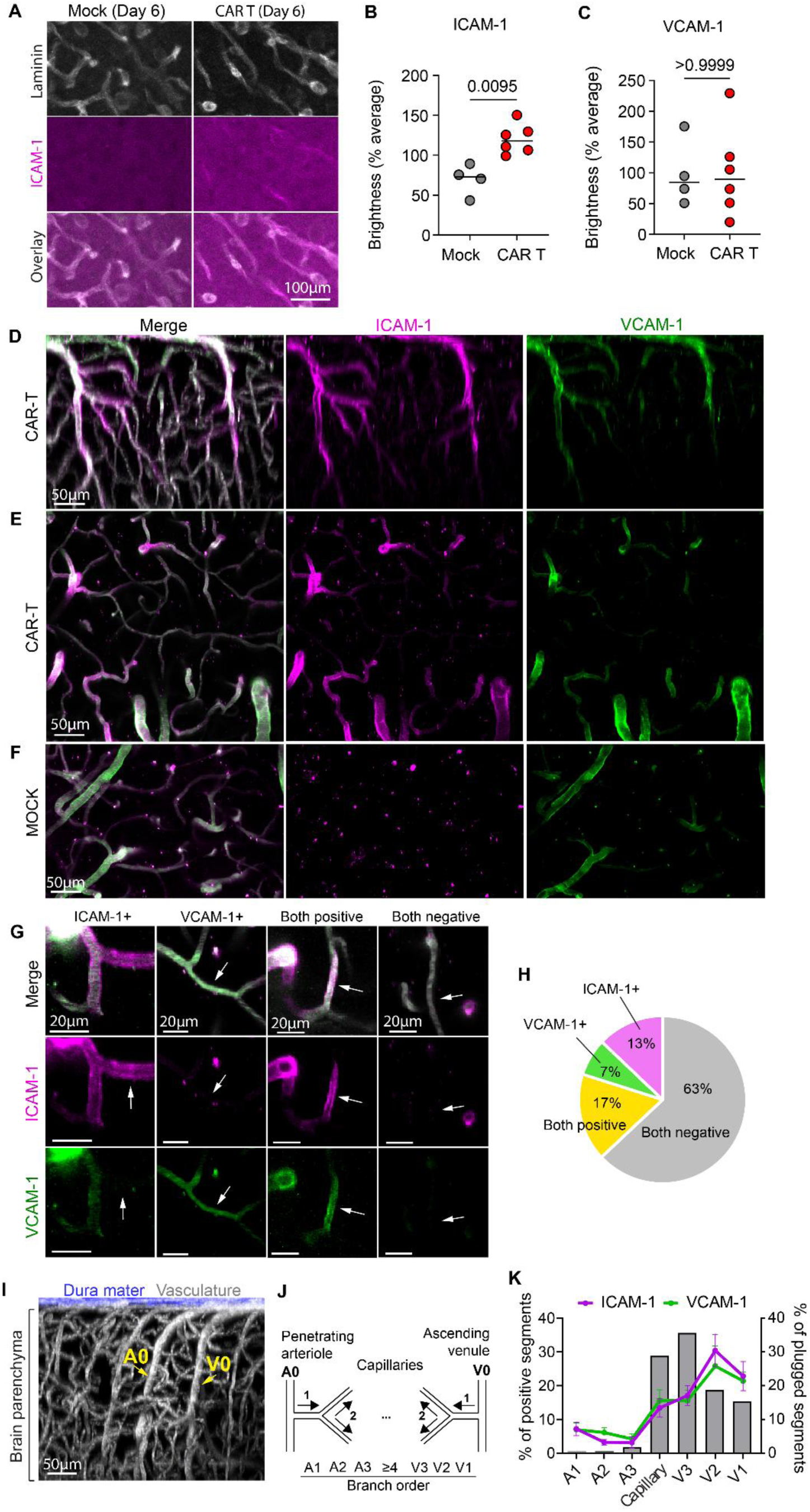
ICAM-1 is widely upregulated in brain capillaries in CD19-CAR T cell treated mice. Wild-type non-cancer bearing mice received lymphodepletion followed by either 10 million CD19-CAR T cells or mock transduced T cells. (**A**) Representative immunofluorescence images on day 6 after CAR T cell treatment, showing brain capillary labeling with anti-laminin antibody (neuronal nuclei are also labeled) and ICAM-1 staining. (**B, C**) Quantification of ICAM-1 and VCAM-1 immunofluorescence brightness, normalized to the average of all mice within each experiment. Each dot indicates one mouse, N=4-6 mice per group, Mann-Whitney test. (**D-F**) Representative in vivo two photon images of the cortical microvasculature on day 5 after treatment with CAR T cells **(D,E)** or mock transduced T cells **(F)**. ICAM-1 and VCAM-1 were labeled simultaneously with i.v. fluorescent antibodies, intravascular dextran label is shown in white for the merged images. **(G)** Representative images of cortical microvascular segments expressing ICAM-1, VCAM-1, both, or neither in a CAR T cell treated mouse. Intravascular dextran label is shown in white for the merged images. **(H)** Distribution of capillary segment ICAM-1 and VCAM-1 expression in CAR T cell treated mice. Averages are shown for 3 mice, 3-4 separate z-stacks per mouse, for a total of 2011 capillary segments. **(I)** Representative image of the cortical microvascular architecture. The dura was visualized by second-harmonic generation, the vasculature was labeled with i.v. dextran. A0, penetrating arteriole; V0, ascending venule. **(J)** Schematic showing branch order assignment for the cerebral capillary bed. **(K)** Capillary plugging (right y-axis, grey bars) and ICAM-1/VCAM-1 expression (left y-axis) predominantly occur in the true capillaries and the capillary-venule transition zone. ICAM-1/VCAM-1 expression is shown as the average of 3 mice with error bars showing SEM. The capillary plugging counts show the total number of leukocyte plugs per branch order (N=7 mice).

We then used *in vivo* two-photon imaging to investigate the 3D spatial distribution of ICAM-1 and VCAM-1 expression across vascular zones. Mice implanted with chronic cranial windows were treated with cyclophosphamide preconditioning followed by 10 million CD19-targeted CAR T cells, and imaged on day 5 after CAR T cell infusion. Immediately prior to imaging, we intravenously injected fluorescently labeled antibodies directed against ICAM-1 and VCAM-1 to visualize the expression of both adhesion molecules on the luminal side of the endothelium (Movie S2). Expression of both adhesion molecules was strong and relatively uniform in the pial arterioles and venules (Fig. S2A). We frequently observed adherent leukocytes in the pial venules (Movie S1), but occasionally they even adhered to the walls of pial arterioles despite the high shear stress (Fig. S2B, Movie S3). The pial vessels traverse the surface of the brain and connect to the penetrating arterioles and draining venules, which also strongly expressed ICAM-1 and VCAM-1 (Fig 2D,E). Control mice treated with cyclophosphamide only had minimal ICAM-1/VCAM-1 expression in these zones (Fig. S2C), while it was somewhat more prominent in mice treated with mock transduced T cells (Fig 2F). The increased expression of ICAM-1 and VCAM-1 in meningeal vessels after CAR T cell treatment shows that these vascular zones are highly reactive to inflammatory stimuli, and likely explains why leukocytes are frequently seen arrested in these vessels.

We next assessed the distribution of ICAM-1 and VCAM-1 expression in the microvessels of the brain parenchyma where we had previously observed capillary plugging (Fig 2G). Adhesion molecule expression was often heterogenous between adjacent capillary segments, likely representing individual microendothelial cells with differential reactivity. In CAR T cell treated mice, an average of 13% of capillaries expressed ICAM-1 only, 7% expressed VCAM-1 only, 13% expressed both, and 63% expressed neither (Fig. 2H). In control mice, 80% (mock) to 90% (cyclophosphamide) of capillaries were negative for both ICAM-1 and VCAM-1 (Fig SA,B). While the overall percentage of ICAM-1+ capillary segments after CAR T cell treatment was similar across individual mice, VCAM-1 was more heterogenous between individuals even though they had all received the same CAR T cell treatment (Fig. S2C,D). These findings align with our findings on histologic analysis, where ICAM-1 expression in capillaries was increased over mock controls (Fig 2B), while VCAM-1 was sometimes higher, and sometimes lower (Fig 2C).

Endothelial cell transcriptional profiles vary along an arterial-capillary-venous continuum in the brain, and expression of proinflammatory response profiles is associated with venous identity (*15, 16*). To determine whether these identities affect adhesion molecule expression and consequently the likelihood of capillary plugging, we analyzed expression of ICAM-1 and VCAM-1 across the arteriovenous continuum within the microvascular bed by in vivo two-photon imaging. We assigned microvascular identities as arteriole-capillary transition zone (capillary segments that are 1, 2, or 3 branch orders removed from a penetrating arteriole), true capillaries (>4 branch orders from both arterioles and venules), and the capillary-venule transition zone (1, 2, or 3 branch orders from the ascending venule (Fig 2I,J). Both ICAM-1 and VCAM-1 were predominantly expressed in the capillaries (branch order ≥4) and the capillary-venule transition zone (V1-V3), and much less in the arteriole-capillary transition zone (A1-A3) (Fig 2K, S3E,F). The distribution of capillary plugs was also predominantly on the venular side of the capillary bed. Of 59 plugging leukocytes where branch order could be determined, all but one were in the capillaries or capillary-venule transition zone (Fig 2K). This supports the conclusion that high ICAM-1 and/or VCAM-1 expression predisposes capillaries to leukocyte arrest. However, the capillary plugs were most frequently seen in the capillary zones most distant from the venule, while the highest ICAM-1 and VCAM-1 expression was seen closer to the venule. This suggests that factors other than ICAM-1 and VCAM-1 expression, such as smaller diameters, may also affect the likelihood of capillary plugging.

Taken together, these results show that ICAM-1 and VCAM-1 upregulation and capillary plugging primarily occur on the venular side of the brain capillary network in response to CAR T cell therapy. While ICAM-1 upregulation is consistently higher in CAR T compared to mock treated animals, VCAM-1 upregulation appears more heterogenous.

### CAR T cells preferentially increase affinity to VCAM-1 rather than ICAM-1

The strength of leukocyte-endothelial interaction is regulated by conformational changes in integrins that occur upon leukocyte activation (*14*). We confirmed that in our mouse model, CAR T cells acquire a predominantly effector/activated phenotype (CD62L^-^ CD44^hi^) (Fig. S4A), with the peak of activation by day 4 and remaining sustained through day 6. The endogenous T cells also became activated, but to a lesser degree and on a slower time course (Fig. S4B). T cells remained predominantly naïve in control mice receiving only lymphodepletion with cyclophosphamide (Fig. S3C). These results predict that integrin activation should be greater in CAR T cells compared to endogenous bystander T cells, and absent in control mice.

The key integrins in neuroinflammation are VLA-4 (the integrin α4β1 heterodimer) which binds VCAM-1, and LFA-1 (the integrin αLβ2 heterodimer), which binds ICAM-1 (*14, 17*). To measure the affinity of leukocyte integrins VLA-4 and LFA-1 to their endothelial binding partners, we incubated fresh whole blood from CD19-CAR T cell treated mice with fluorescently labeled ICAM-1 and VCAM-1 peptide multimers (Fig. 3, A and B) and quantified multimer binding to leukocytes by flow cytometry (gating strategy shown in Fig. S5). Somewhat surprisingly, the affinity of T cell LFA-1 to ICAM-1 was not higher in CAR T cell treated mice compared to mock controls on days 5, 12, or 19 after CAR T cell infusion (Fig. 3C). In fact, CAR T cells had lower ICAM-1 affinity than endogenous, CAR-negative circulating T cells on day 5 and 12 after CAR T cell infusion (Fig. 3C, S6A,B). Although cell surface expression of integrin αL (LFA-1) in T cells was higher in CAR T cell treated mice compared to mock controls (Fig. 3D), its expression actually dropped in CAR T cells over time, whereas endogenous T cells in the same mice saw a transient increase (Fig. 3D, S6E,F). The pattern of decreasing LFA-1 expression and affinity over time does not match the kinetics of T cell activation and capillary leukocyte adhesion, which peaks during days 4-6 after CAR T cell infusion. Thus, even though ICAM-1 is broadly upregulated after CAR T cell treatment across the brain endothelium in vivo, on the T cell side the ICAM-1/LFA-1 interaction appears to be deemphasized.

**Fig. 3.**
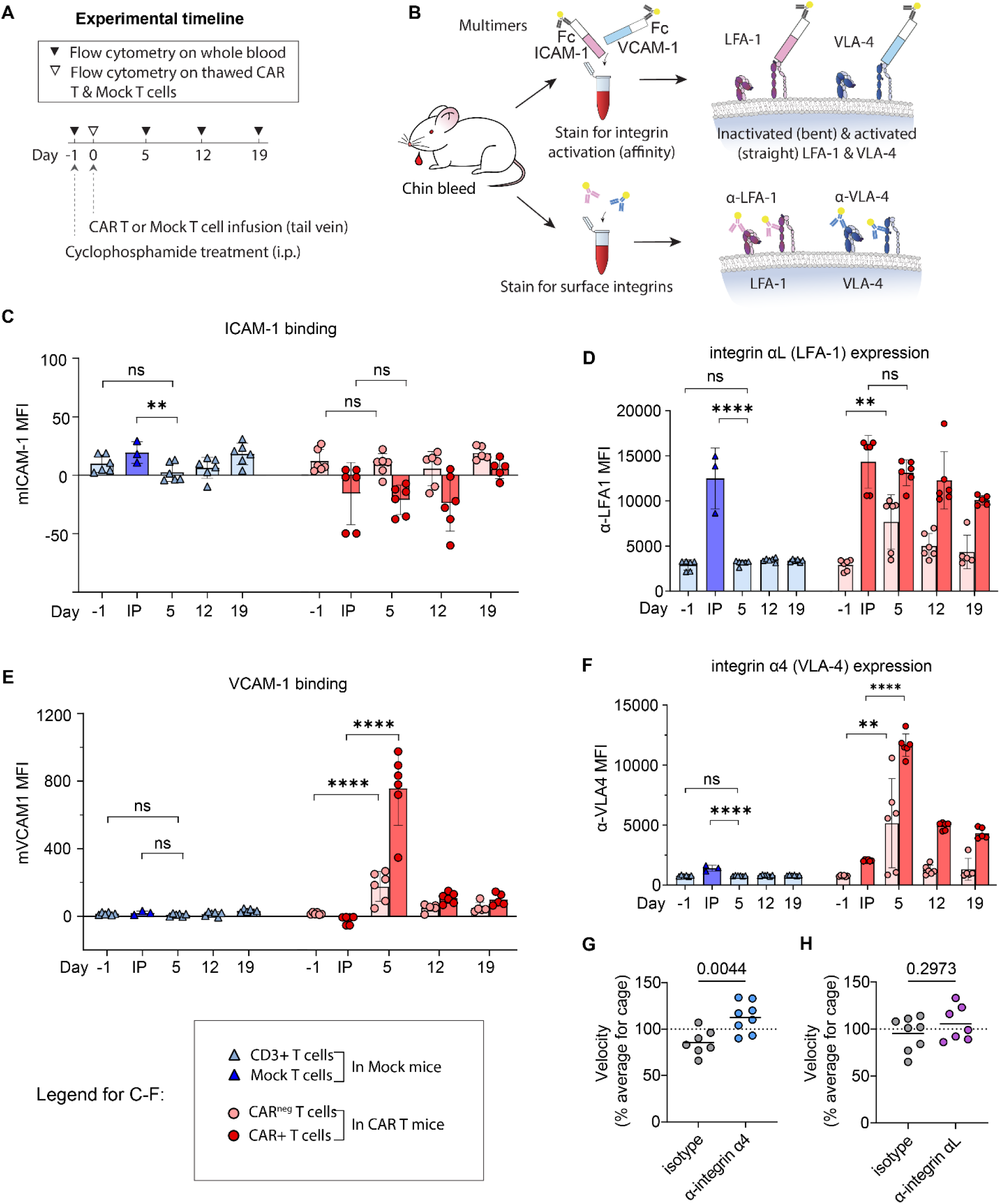
CAR T cells strongly upregulate affinity of VLA-4 to VCAM after infusion. (**A**) Experimental timeline. Mice received 10 million CD19-targeting CAR T cells or mock transduced T cells on day 0, and blood samples were taken on days 5, 12 and 19 to assess integrin activity on circulating immune cells. (**B**) Whole blood was labeled with either ICAM-1/VCAM-1 multimers tagged with anti-human Fc antibodies, or with antibodies that label integrins regardless of conformation, and quantified by flow cytometry. (**C-F**) Quantification of flow cytometric measurements of T cell affinity to ICAM-1 (C), cell surface expression of integrin αL (LFA-1, D), affinity to VCAM-1 (E), and surface expression of integrin α4 (VLA-4, F). The y-axis indicates the mean fluorescence intensity (MFI, arbitrary units) for each analyte, and the x-axis indicates time relative to CAR T cell infusion, IP = cryopreserved infusion product immediately upon thawing, prior to infusion into mice. In mice treated with mock transduced T cells (blue triangles), data show either circulating T cells (combination of mock and endogenous) or the cryopreserved mock T cells (IP). For mice treated with CD19-CAR T cells (red circles), data show circulating Thy1.1+ CAR+ T cells, Thy1.1- endogenous CAR^neg^ T cells, or cryopreserved CAR T cells (IP). Each data point indicates one mouse, the box plots show the mean and the whiskers show the standard deviation. N=3 independent experiments, for a total of 3-6 mice per group, one-way ANOVA with Dunnett’s multiple comparisons posttest, only comparisons between day -1 or IP to day 5 are shown for clarity. *P<0.05, **P<0.01, ***P<0.001, ****P<0.0001, ns P>0.05. (**G, H**) Scatter plots showing movement velocity in the open field on day 6 after CAR T cells. Mice were treated with antibodies against integrin α4 (G) or integrin αL (H), or isotype control, on days 1, 3, and 5. Values are normalized to the average for each cage of mice. N=7-8 per group in 7 independent experiments, each dot indicates one mouse, unpaired 2-sided t test.

In contrast, T cell expression of integrin α4 and the affinity of VLA-4 to VCAM-1 increased dramatically after CAR T cell infusion (Fig. 3E,F). Both CAR T cells and endogenous T cells showed the same behavior, but it was much more pronounced in CAR T cells. Both subsets of T cells showed a sharp increase in VCAM-1 binding by day 5, followed by a decline without return to baseline by day 12 and 19 (Fig. S6C,D). For CAR T cells, both VCAM-1 binding and integrin α4 (VLA-4) expression rose more strongly in the CD8+ subset (Fig. S7D,H). In contrast, among endogenous T cells, CD4+ T cells had higher ICAM-1 and VCAM-1 affinity than CD8+ T cells (Fig. S7A,C), even though CD8+ T cells more strongly upregulated integrin expression (Fig. S7E,G). This shows how integrin expression and integrin affinity can be uncoupled depending on the specific activation state of the T cell.

Taken together, these results indicate that CAR T cell binding to VCAM-1 but not ICAM-1 is activated when the CAR T cells encounter antigen in vivo. The concomitant increase in VLA-4 affinity of endogenous T cells may be mediated by cell-cell signaling, and/or by the proinflammatory cytokine/chemokine milieu (*14*).

We also measured the affinity of LFA-1 and VLA-4, as well as the surface expression of integrins αL and α4, on Ly6G+ neutrophils and CD11b+/Ly6G- monocytes/macrophages. In both cell populations, we observed no significant changes in integrin affinity between CAR T treated mice and mock controls (Fig. S8A-D). Similarly, surface expression of integrins αL and α4 did not change, with the exception of a slight increase in VLA-4 expression on CD11b+/Ly6G- myeloid cells in CAR T mice on day 5 (Fig. S8E-H). These results show that increases in integrin activation and expression are largely restricted to T cells, and most prominent in CAR T cells.

Our data so far suggests that the interaction of VLA-4 with VCAM-1 is strongly modulated during CAR T cell treatment, and its dynamics align with the timing of CAR T cell neurotoxicity, with a peak on day 5 after CAR T cell infusion. We reasoned that blockade of this interaction may decrease clinical CAR T cell neurotoxicity. We have previously shown that locomotion activity is severely impaired in mice experiencing neurotoxicity after CD19-CAR T cell treatment (*5*). We treated mice i.p. with 2mg/kg per dose of antibodies directed against integrin α4 or integrin αL on days 1, 3, and 5 after CAR T cell infusion and measured spontaneous locomotion in an open field. Mice receiving integrin α4 blockade covered a median of 27% more distance compared to mice receiving isotype control antibody (P=0.0044, n=7-8 mice per group) (Fig. 3G), whereas no significant effect was seen with integrin αL blockade (P=0.2973) (Fig. 3H). This data again supports the conclusion that the VLA-4/VCAM-1 interaction plays a more important role than the LFA-1/ICAM-1 interaction in mediating CAR T toxicity.

### Cytokines induce ICAM-1 and VCAM-1 expression in human brain endothelium in vitro

CAR T cell treatment can lead to extremely high cytokine levels in the blood, and elevations of multiple blood cytokines have been associated specifically with ICANS in clinical studies (*4*). We have also shown similar increases in IL-6, IFN-γ, and other serum cytokines in our mouse model (*5*). It follows that these cytokines in the circulation might be sufficient to increase expression of ICAM-1 and VCAM-1 in the brain microendothelium. Proinflammatory cytokines, including TNF, IL1-β and IFN-γ, are known to activate endothelial cells and upregulate adhesion molecule expression (*18*). However, most prior studies have used very high, supraphysiologic doses of cytokines applied in vitro to endothelial cells, and the brain-specific microendothelial responses have not been well characterized. To determine the effect of ICANS-associated cytokines on the brain microendothelium, we applied a panel of cytokines to primary human brain microendothelial cell (HBMEC) monolayer cultures and measured ICAM-1 and VCAM-1 expression by flow cytometry. The cytokine concentrations were chosen to reflect blood levels in CAR T cell patients during CRS and/or ICANS (*3, 19–22*), reviewed in Gust et al. (*4*). Healthy control blood cytokine levels were per Bender et al., Kim et al., and Yan et al. (*12, 23–25*). Of the cytokines we tested, only TNF, IL-1β and IFN-γ induced ICAM-1 (Fig. 4A). VCAM-1 expression was more weakly upregulated by the same concentrations of TNF and IL-1β. The other cytokines, including IL-6, had no effect on ICAM-1 or VCAM-1 expression (Fig. 4A, S9). These results confirm that brain microendothelial cells upregulate ICAM-1 even with low levels of proinflammatory cytokine stimulation, whereas VCAM-1 responds less vigorously. This correlates reasonably well with our *in vivo* mouse data, where ICAM-1 expression rose more strongly and consistently than VCAM-1 expression in CAR T treated mice. However, our cultured primary HBMECs likely represent a mix of cells from multiple vascular zones, and specialized capillary-venule transition zone cells with higher VCAM-1 expression may be underrepresented.

**Figure 4.**
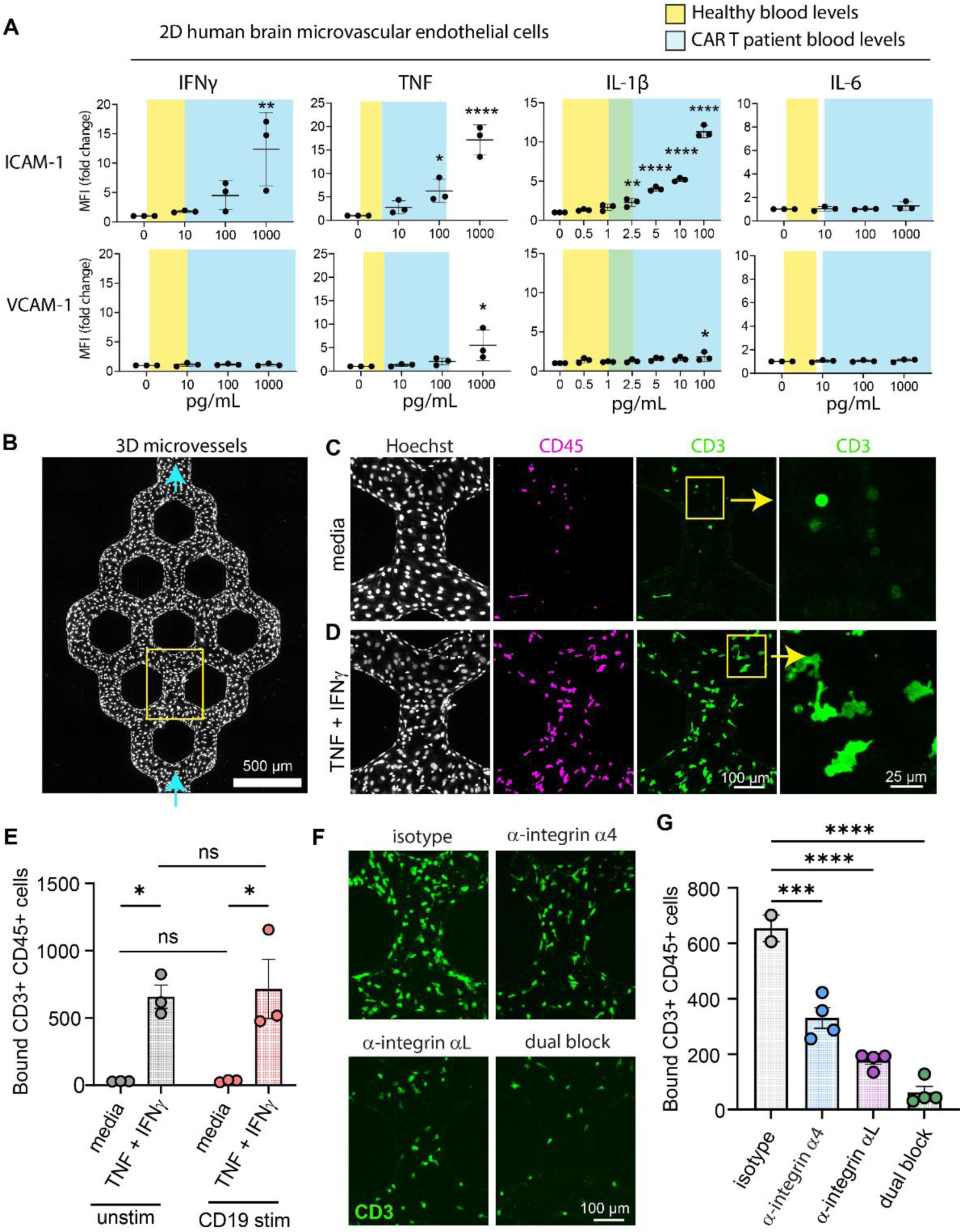
Cytokines induce ICAM-1 and VCAM-1 expression and CAR T cell adhesion on human brain microendothelium. (**A**) ICAM-1 and VCAM-1 expression were measured by flow cytometry in HBMEC monolayer cultures after treatment with cytokines. The x-axis indicates the concentration of each cytokine that was added to the culture media for 24h. The y-axis shows the fold change in mean fluorescence intensity compared to the 0 pg/mL vehicle-only control. Y-axes are the same for ICAM-1 and VCAM-1 for each cytokine to allow comparison of effect size. The yellow shading indicates the approximate concentrations of each cytokine found in the blood of healthy volunteers, the blue shading indicates the range of concentrations reported in CAR T cell patients, and green shows overlap of these ranges. N=3 independent experiments, one-way ANOVA with Dunnett’s multiple comparisons posttest. Only comparisons with the 0 pg/mL vehicle-only control were performed, P>0.05 not shown for clarity. (**B**) Representative image of a 3D HBMEC microvessel that was fixed and immunolabeled after perfusion with CAR T cells. The arrows indicate the direction of flow. The yellow box indicates the area magnified in (C) and (D). Representative examples are shown of a media-only pretreated vessel (**C**) and a vessel pretreated with 100pg/mL each of TNF and IFNg (**D**) prior to perfusion with CAR T cells. CAR T cells are immunolabeled for CD45 (magenta) and CD3 (green). The nuclei of endothelial cells and CAR T cells are labeled with Hoechst 33342. The yellow boxes show the area magnified in the last column to show the difference in adherent CAR T cell morphology between conditions. (**E**) Quantification of adherent CD3+ CD45+ CAR T cells per microvessel after 1h of perfusion followed by 2 media washes. Prior to perfusion, CAR T cells were either rested on uncoated plates (“unstim”) or stimulated with plate-bound human CD19 peptide (“CD19 stim”). Vessels were either pretreated with media only, or with 100pg/mL each of TNF and IFN-γ. N=3 microvessels per condition, 2-way ANOVA with Tukey’s multiple comparisons posttest. (**F**) Representative examples of microvessels treated with integrin blockade, images are magnifications of the area indicated by the yellow box in (B). All vessels were pretreated with 100pg/mL each of TNF and IFN-γ, and all CAR T cells were stimulated with plate-bound CD19. During CAR T cell perfusion, 50μg/mL of natalizumab (anti-integrin α4), TS1/22 (anti-integrin αL), or both were added to the media. Adherent CAR T cells are shown by CD3 immunolabeling (green). (**G**) Quantification of adherent CAR T cells in the experiment described in (F). N=2-4 microvessels per condition, one-way ANOVA with Dunnett’s multiple comparisons test, only comparisons with media control were performed. **For all graphs:** *P<0.05, **P<0.01, ***P<0.001, ****P<0.0001, ns P>0.05.

### CAR T cell adhesion in 3D microvessels is dependent on both LFA-1/ICAM-1 and VLA-4/VCAM-1 interactions

To model the interaction of the brain microendothelium with CAR T cells in a fully human system, we created collagen 3D patterned microvessels that are endothelialized with primary HBMECs (*26*)(Fig. 4B). We then perfused human CD19-CAR T cells through these vessels, and measured their ability to adhere to the endothelial wall. In vessels not exposed to cytokines, only few CAR T cells adhered, and those that did remained predominantly rounded in shape (Fig. 4C). When vessels were pretreated with 100pg/mL of each TNF and IFN-γ to mimic the cytokine environment in patients, CAR T cells adhered in large numbers and assumed an ameboid morphology (Fig. 4D). Interestingly, stimulating the CAR T cells with plate-bound CD19 prior to perfusion was not sufficient to induce adhesion without cytokine pretreatment of the vessel (Fig. 4E). We ensured that CAR T cells did not receive any direct cytokine stimulation by washing the vessels with cytokine-free media prior to CAR T cell perfusion. This indicates that activation of the endothelium alone is sufficient to induce vigorous T cell adhesion, and that simply encountering a CAR target is not sufficient if the endothelium is not activated. We then tested whether the CAR T cell-endothelial interaction could be disrupted by clinically relevant antibodies against VLA-4 (natalizumab)(*27*) and/or LFA-1 (clone TS1/22). We chose TS1/22 because it binds the same integrin αL domain as efalizumab (*28*) but does not cross-react with VLA-4 (*29*). Antibody concentrations were chosen to represent typical blood levels in patients (*30, 31*). Both VLA-4 and LFA-1 blockade decreased CAR T cell binding, but LFA-1 was more effective (Fig. 4F,G). When both antibodies were combined, CAR T cell binding was almost completely abolished (Fig 4F,G). Together, these results show that human brain microendothelial activation by cytokines upregulates ICAM-1 and VCAM-1, which are necessary and sufficient to synergistically mediate CAR T cell adhesion to the vessel wall. The 3D model suggests that adhesion to the vessel wall can occur without prior T cell stimulation, at least at the high end of the range of cytokine release that occurs after CAR T cell treatment.

### In CD19-CAR T cell patients, increases in soluble ICAM-1 and VCAM-1 are associated with ICANS and correlate with serum cytokine levels

Next, we sought to determine if ICAM-1 and/or VCAM-1 levels are also associated with ICANS in human CAR T cell patients. It would stand to reason that the increased cytokine levels that are seen after CD19-CAR T cell infusion (*4*) can induce ICAM-1 and VCAM-1 expression in the endothelium, similar to what we found in vitro in brain microendothelial cells and in the ICANS mouse model. We measured the concentration of soluble ICAM-1 and VCAM-1 in the plasma of CD19-CAR T cell treated patients at baseline (day 1 after CAR cell infusion) and on day 7 after CAR T cell infusion in 3 different clinical trial cohorts of patients with acute lymphoblastic leukemia. This included adults treated with CD19-directed CAR T cells (n=48 patients, 50% with ICANS) (*32*), children and young adults treated with CD19-CAR (n=37 patients, 49% with ICANS) (*33, 34*), and children and young adults treated with CD19/CD22-CAR (n=24 patients, 33% with ICANS)(*35*). In all cohorts, the ratio of day 7 to day 1 soluble ICAM-1 and VCAM-1 levels was higher in patients with ICANS compared to patients without ICANS, but this difference was only statistically significant in the adult cohort for ICAM-1 (Fig. 5A, Table S1). Expressing the rise as the absolute difference between day 7 and day 1 provided similar results (Table S1). The two patients with fatal (grade 5) ICANS had some of the highest ratios of day 7/1 ICAM-1 (4.6, 3.6) and VCAM-1 (2.6, 2.2). Baseline ICAM-1 and VCAM-1 levels were not different between patients with or without subsequent ICANS (Table S1). We next determined whether the cytokines that induced ICAM-1 and VCAM-1 in vitro also correlated with in vivo induction in patients. Indeed, higher serum levels of IFN-γ and TNF on day 7 after CAR T cell infusion were associated with greater rises in soluble ICAM-1 and VCAM-1 (Fig. 5 B,C). Although IL-6 did not induce either adhesion molecule in vitro, patients with serum IL-6 levels of >100pg/mL also had a significantly greater rise in soluble ICAM-1. This data must be interpreted with the caveat that patients with high IL-6 levels also typically have high IFN-γ and TNF levels. In addition, the measurements of soluble ICAM-1 and VCAM-1 reflect the systemic circulating molecules and are not specific to the brain endothelium. Nonetheless, this data is consistent with a mechanism of toxicity where increased ICAM-1 and/or VCAM-1 expression in the brain endothelium is induced by circulating proinflammatory cytokines.

**Fig. 5.**
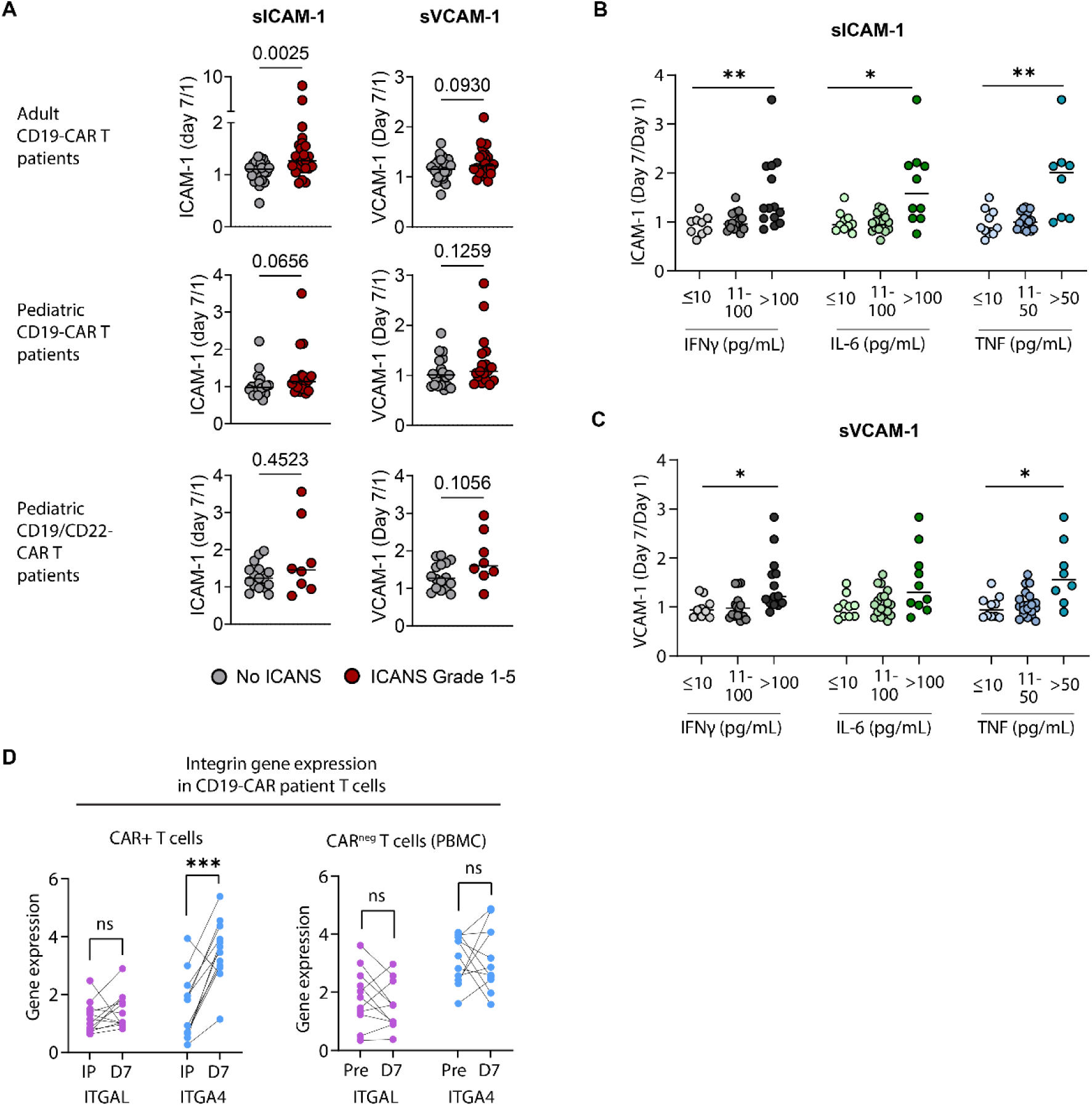
Endothelial-leukocyte adhesion molecule dynamics in CD19-CAR T cell patients. (**A**) Patients with ICANS had greater rises in soluble ICAM-1 or VCAM-1 in three independent clinical cohorts. The ratio of day 7/day 1 serum levels is shown for each patient, the line shows the median, Mann-Whitney test. (**B, C**) Patients with the highest day 7 serum levels of TNF, IFN-γ and IL-6 also have the greatest increase in soluble ICAM-1 and VCAM-1 (Day 7/Day 1 ratio). Patients were grouped by cytokine levels analogous to the concentrations used in the in vitro experiments in Figure 4. The lines indicate the median, Kruskal-Wallis test with Dunn’s multiple comparisons posttest, only comparisons with the <10pg/mL group were performed. *P<0.05, **P<0.01, all other comparisons were P>0.05. (**D**) Integrin α4 expression increases in CAR T cells after infusion. Integrin gene expression levels are shown for sorted CAR T cells or CAR-negative T cells by pseudobulk reanalysis of single cell RNA sequencing data. Each data point represents one sample, paired samples from individual patients are connected by lines. IP, infusion product; D7, PBMCs on day 7 after CAR T cell infusion; Pre, baseline PBMC sample prior to CAR T cell infusion. Paired 2-sided t test, *P<0.05, **P<0.01, ***P<0.001, ****P<0.0001, ns P>0.05.

### CAR T cells upregulate integrin α4 expression in human CD19-CAR T cell patients

Our mouse data indicated that the interaction of VLA-4 with VCAM-1 is strongly upregulated in vivo by CAR T cells and endogenous T cells. To determine if this is the case in human CAR T cell patients, we reanalyzed a published single-cell RNA sequencing dataset from patients treated with commercially available CD19-CAR T products (*36*). We compared pseudobulk gene expression levels of integrin α4 (ITGA4) and integrin αL (ITGAL) between CAR T cells in the infusion product and CAR T cells sorted from day 7 PBMCs (n=12 patients), and between circulating non-CAR T cells at baseline and day 7 (n=11 patients). Similar to what was seen in mouse CAR T cells, ITGA4 expression rose substantially in CAR T cells after they were infused (P=0.0002) while ITGAL expression did not change (P=0.2874) (Fig. 5D). Endogenous T cells did not see a concomitant change in either integrin’s expression level between baseline and day 7 after CAR T cell infusion (Fig. 5D). This is unlike in the mouse model, where both increased. This difference could be due to the fact that the mice were healthy before CAR T cell treatment, whereas the patients may have had previously activated circulating T cells due to multiple prior therapies, lymphodepleting chemotherapy, and/or infections.

## DISCUSSION

This study supports a model of ICANS as a neurovascular syndrome, where brain capillaries are plugged by circulating leukocytes in response to increased leukocyte-endothelial affinity. The resultant brain tissue hypoxia could be sufficient to cause transient cognitive dysfunction, which would be expected to resolve once the plugging is cleared. This is consistent with the natural history of ICANS in cancer patients, where symptoms typically reach their maximum within a few days after onset, and then gradually resolve (*3, 19*). If plugging were to become very severe, it could lead to perfusion failure, overt tissue injury, and even cerebral edema due to breakdown of the blood-brain barrier.

Clinically, the risk of ICANS is strongly associated with preceding CRS, and ICANS typically develops soon after the peak of CRS (*4*). This observation led us to the straightforward hypothesis that capillary plugging is induced by cytokine release with concomitant endothelial activation and leukocyte activation during the acute inflammatory phase of CAR T cell therapy. Such a mechanism would be independent of any off-tumor on-target activity of the CAR T cells and would be consistent with ICANS as an overarching syndrome of neurotoxicity that could occur with any cytokine-activating therapy or pathology.

The findings in this study support the hypothesis of cytokine increase leading to endothelial and leukocyte activation, followed by slowing and sticking of leukocytes in tiny brain vessels. Our use of a fully murine in vivo model is key for this research, because even mice with humanized immune systems likely cannot recapitulate the full spectrum of cell-cell interactions in the brain neurovascular unit. In human in vitro systems, we show that the brain endothelium can be activated by relatively modest increases in IL-1β, IFN-γ and TNF, reflective of blood levels that are typically found in CAR T cell patients. This is sufficient to induce vigorous adhesion of CAR T cells to the endothelium, which is prevented by VLA-4 and/or LFA-1 blockade. In human patients, we find an association of IFN-γ and TNF levels with increased soluble ICAM-1 and VCAM-1, again consistent with our hypothesis. These findings lead to further complex questions, as currently there are no reliable techniques to directly prove or disprove in human patients whether brain capillary plugging causes ICANS. It remains unknown why ICANS is such a brain-specific syndrome. If endothelial-leukocyte adhesion was broadly upregulated throughout the body, one would expect predictable organ dysfunction in other organs such as the kidneys. However, this has not been observed.

Our results suggest a differential contribution of ICAM-1 and VCAM-1 interacting with their respective leukocyte integrins, and many questions remain regarding the regulation and consequences of these differences. In our mouse model, we show that circulating CAR T cells and endogenous T cells increase their affinity to VCAM-1 but not ICAM-1, and only blockade of the VLA-4/VCAM-1 but not the LFA-1/ICAM-1 interaction leads to behavioral improvement. Similarly, only VLA-4 but not LFA-1 expression increases in CAR T cells in human patients. On the endothelial side however, ICAM-1 appears to be somewhat more strongly upregulated than VCAM-1. In vitro, brain microendothelial cells upregulated ICAM-1 more strongly than VCAM-1 in response to cytokine stimulation. Our in vivo imaging studies show that both ICAM-1 and VCAM-1 increase on the venular side of the brain capillary network, which is also where the majority of the leukocyte plugging occurs. Most of the activated capillary segments expressed either ICAM-1 or both ICAM-1 and VCAM-1, while fewer only had VCAM-1. One possible explanation is that increased VLA-4 affinity is sufficient to induce leukocyte adhesion to the endothelium even if VCAM-1 is only expressed at basal or slightly increased levels. To better understand the contribution of the ICAM-1/LFA-1 vs. the VCAM-1/VLA1-1 axis in human patients, future studies should measure integrin affinity/avidity in freshly isolated patient CAR T cells and other circulating leukocytes.

Could our data also be consistent with alternative models of ICANS? The function of the ICAM-1/LFA-1 and VCAM-1/VLA-4 axes extends well beyond the leukocyte-endothelial interaction. They are also important for cell-cell signaling in the immune system, immune synapse formation, and immune cell trafficking in tissues (*14*). Thus, other potential mechanisms of CAR toxicity should also be considered, such as dysfunctional activation of the myeloid compartment (*37*), or direct tissue invasion. However, tissue invasion or myeloid activation on their own would not necessarily cause brain dysfunction, and another step in the injury mechanism would be required. One such mechanism could be off-tumor, on-target activity, where CAR T cells destroy normal bystander cells that express the target antigen. Although there is low level expression of CD19 in some cell types in the fetal human brain (*38*), there is no convincing evidence to date that it is present in the mature brain. Contrast this with CD22, which is also commonly targeted with CAR T cells and is expressed by oligodendrocytes but has not been associated with high neurotoxicity (*39*). A structural injury would also not be expected to be so quickly reversible. ICANS-associated cytokines have many other direct effects on the brain (*4*), including loss of endothelial tight junction integrity, and activation of other cell types such as microglia (*37, 40*). Even if these other processes are also present during ICANS, increased endothelial and/or T cell activation and subsequent capillary plugging alone should be sufficient to induce hypoperfusion and cognitive dysfunction. In vivo oxygen imaging studies in mouse show that oxygen delivery drops rapidly when individual capillaries stall (*12*). Modeling of network effects of capillary stalls shows that even small increases in capillary stalling lead to exponentially more severe failure of brain perfusion (*41, 42*). Mouse studies show that capillary plugging may also contribute to brain dysfunction during sepsis, seizures, and neurodegenerative disorders (*6–8*).

If capillary plugging is indeed responsible for at least aspects of neurotoxicity, what are the therapeutic or preventive options? Blockade of ICAM-1, integrin α4, or IL-1β has been previously tested in patients with ischemic stroke, after mouse studies suggested that decreasing leukocyte recruitment to ischemic vasculature may improve outcomes (*43*). This approach has been all but abandoned after high quality clinical trials showed lack of efficacy or even adverse outcomes (*43*). In addition, the anti-integrin antibodies natalizumab and efalizumab have both been associated with increased rates of progressive multifocal leukencephalopathy due to CNS-specific immunosuppression, although this only occurs after prolonged exposure (*44, 45*). Use of IL-1β blockade to prevent ICANS has been supported by a mouse study (*26*), which however did not investigate the possible mechanism of injury that might be mediated by IL-1β. Nonetheless, IL-1β blockade with anakinra has become a mainstay of ICANS treatment. This is despite a paucity of data reporting IL-1β levels in the blood or CSF of CAR T patients, and several clinical studies that failed to find a protective effect of anakinra prophylaxis against ICANS (*46, 47*). The most promising approach may be to engineer CAR T cells to reduce their potential for setting off dysfunctional inflammatory cascades. This could be done by co-expressing cytokine blocking antibodies (*48*), or inhibiting integrin expression (*49*) to reduce neurotoxicity while preserving antitumor activity.

In conclusion, our study shows that the systemic cytokine release after CAR T cell treatment can be sufficient to cause brain endothelial activation and increased adhesion molecule expression. At the same time, integrin affinity increases not only in CAR T cells, but also in endogenous T cells. The relative contribution of T cell integrin activation and increased endothelial adhesion molecule expression remains to be further studied.

## MATERIALS AND METHODS

### Study design

Sample size was prespecified for the mouse *in vivo* experiments by power analysis to achieve 80% power at an alpha level of 0.05. Effect size was estimated by pilot experiments.

To avoid bias, treatment was randomly assigned to animals after baseline imaging and behavioral testing. All image and data analysis was blinded to group allocation and treatment conditions.

No outliers were excluded, and all experimental results were included in the analyses.

### Animals

5-10 week old wild-type BALB/c mice were used for all experiments. Mice were obtained from Jackson Labs (Bar Harbor, ME) and housed in specific pathogen free facilities. Male and female mice were used in equal numbers, and sexes were randomly assigned to experiments. We have previously shown that there is no sex difference in CAR T neurotoxicity in human patients (*3, 19*) or in our mouse model (*5*). Since CAR T treatment related toxicity manifestations vary across cohorts of mice and batches of CAR T cells, mouse data was always compared to cagemate controls and, where appropriate, normalized by average values within a cage.

### Murine CAR T cell generation

Our murine CD19-directed CAR consists of an anti-murine-CD19 single chain variable fragment (scFv) derived from the 1D3 hybridoma (*50*), a CD28 transmembrane domain and murine intracellular 4-1BB and CD3z costimulatory domains. It is followed by a T2A sequence and Thy1.1 or tdTomato as a transduction marker. The construct was cloned into a MMLV backbone, and gammaretrovirus was produced by transient calcium phosphate transfection of a Phoenix-ECO producer line (ATCC). CAR T cells were generated per published protocol (*51*). Briefly, T cells were collected by mechanically dissociating spleens and lymph nodes of wild type BALB/c mice, followed by MACS selection with CD90.2 beads (Miltenyi Biotec). T cells were then stimulated with CD3/CD28 Dynabeads (ThermoFisher) (days 1-4), murine IL-2 (Peprotech) (days 1-3) or murine IL-15 (Peprotech) (day 4) in RPMI supplemented with 10% fetal bovine serum, penicillin/streptomycin, β-mercaptoethanol, and sodium pyruvate. On days 2 and 3, cells were spinoculated with viral supernatant at 2000xg for 1 hour on retronectin (Takara) coated plates. Transduction efficiency was verified by flow cytometry for Thy1.1. Mock transduced T cells underwent the same transduction protocol, while replacing the viral supernatant with media during the spinoculation step.

### CAR T cell treatment

All CAR T cell treatments were conducted in wild type, non-tumor bearing mice. To reduce rejection of CAR T cells, mice received lymphodepleting chemotherapy with 250mg/kg cyclophosphamide via a single i.p. injection (Day -1). 24h later (Day 0), 10 x 10^6^ CAR T cells or mock transduced T cells per animal were infused via the lateral tail vein. Mice were monitored with daily weight and temperature measurements, and both controls and CAR T treated animals received daily subcutaneous normal saline boluses (20mL/kg). Blood for analysis was collected via chin bleed, retro-orbital puncture under sedation, or post-euthanasia.

### Cranial window preparation and anesthesia

We created thinned-skull windows or chronic skull-removed cranial windows over the somatosensory cortex as previously described (*52, 53*). Briefly, anesthesia was induced with 4% isoflurane and additional pain control provided with buprenorphine. Mice were maintained at 1-2% isoflurane for surgery. After removing the skin, a custom head mount was attached using dental cement. For thinned-skull windows, the skull was thinned to translucency with a handheld drill and a 4mm coverslip was mounted on top of the area with cryanoacrylate instant adhesive. Imaging was started 1 day after the surgery. For chronic windows, the skull was removed, keeping the dura intact, and a plug fashioned of a 3mm and 4mm coverslip glued together with cryanoacrylate adhesive was mounted into the skull defect. Mice were allowed to recover for 2-3 weeks prior to imaging. During imaging, mice were anesthetized with 1.5% isoflurane and mounted in a custom head holder.

### In vivo two photon imaging

Imaging was performed on a Bruker Investigator microscope with Prairie View imaging software, coupled to a Spectra-Physics InSight X3 tunable laser (680-1300nm). Green, red, and far-red fluorescence emission was collected through 525/70nm, 595/50nm, and 660/40 bandpass filters, respectively. Images were collected using a 20X (1.0 NA) water-immersion objective (Olympus; XLUMPLFLN).

All antibodies and intravascular tracers were delivered by injection into the retro-orbital sinus. The blood plasma was labeled with 2mDa dextran conjugated to either FITC or Alexa 680 (2.5-5% in PBS, 20 µl once). The following monoclonal antibodies conjugated to either FITC, Alexa 488, Alexa 568, or Alexa 647 were injected and allowed to equilibrate for at least 15 minutes prior to imaging: anti-CD45.2 (clone 104, Biolegend; 0.4mg/kg per dose), anti-ICAM-1 (clone YN1/1.7.4, Biolegend, cat # 116102, 1mg/kg per dose), and anti-VCAM-1 (clone 429, Biolegend, cat # 105705, 1mg/kg/dose).

### In vivo ICAM-1 and VCAM-1 quantification

In vivo two-photon imaging for ICAM-1 and VCAM-1 measurement was performed on 5 post-CAR T or mock T cell treatment, or on day 6 after cyclophosphamide for cyclophosphamide-only controls. Fluorescence-conjugated monoclonal antibodies against ICAM-1 and VCAM-1 were administered 30 minutes to 1 hour prior to in vivo two-photon imaging. Either of two fluorescence labeling combinations were used: (1) FITC-conjugated ICAM-1 Ab and Alexa 594-conjugated VCAM-1 Ab, together with Alexa 680-conjugated dextran, or (2) Alexa 594-conjugated VCAM-1 Ab and Alexa 647-conjugated ICAM-1 Ab, together with FITC-conjugated dextran. In both cases, an 800nm laser wavelength was used to excite FITC and Alexa 594 simultaneously. 1100nm excitation was used for Alexa 680, and 1200nm for Alexa 647. Three or four individual imaging fields of view, each approximately 289-385 μm (x) x 289-385 μm (y) x 300 μm (z-depth, from the dural surface) were acquired per mouse within each cranial window.

All image processing was performed using Fiji (ImageJ, NIH, US). Within each imaging spot, the total number of microvessel segments was quantified, excluding penetrating vessels. Vessel segments meeting both morphological criteria – elevated fluorescence intensity at the luminal surface of the endothelium and discrete endothelial cell-scale expression patterns – were designated as reference segments. Fluorescence intensity values were measured for these reference segments within each imaging field of view, and their mean intensity was used as the local threshold for identifying additional ICAM-1 and/or VCAM-1-positive segments in the same imaging field. The branch order of each ICAM-1 and/or VCAM-1 positive segment was determined by tracing vessel segments either from the nearest penetrating arteriole or ascending venule (designated as 0^th^ branch order), with the count incremented by 1 at each subsequent bifurcation. Pial arterioles and venules were identified based on their morphological features. Arterioles were characterized by a relatively linear trajectory with gradual reduction in caliber upon bifurcation, whereas venules are more tortuous with frequent branches of variable diameters. Additionally, pial arterioles are located above pial venules. Vessel segments at four or more branching orders distal to the nearest penetrating arteriole or ascending venule were classified as true capillaries.

### Hypoxyprobe quantification

Pimonidazole hydrochloride (Hypoxyprobe Inc. HP3-100Kit) was freshly prepared at 60µg/ml in sterile-filtered PBS and retro-orbitally injected at a concentration of 60 mg/kg under isoflurane anesthesia. 90 minutes after hypoxyprobe injection, the mice were deeply anesthetized and transcardially perfused with PBS followed by 4% PFA. Brains were extracted and immersed in 4% PFA for 24 hours, then transferred to 15% (w/v) and 30% (w/v) sucrose for cryoprotection on subsequent days. Brains were embedded in OCT compound and cut into 50μm coronal sections. Sections were incubated with anti-pimonidazole antibody (Pab2627 rabbit antiserum; 1:100; Hypoxyprobe, Inc.; HP3-100Kit) in antibody solution (10% normal donkey serum, 2% Triton X-100, 0.1% sodium azide in PBS) overnight at room temperature. The following day, sections were incubated with fluorescently labeled secondary antibodies (goat anti-Rabbit IgG Alexa Fluor™ 488, Thermo Fisher) for 2 hours at room temperature. Stained sections were imaged on an Olympus VS120 slide scanner using identical laser and detector settings for each slide. For quantification, 2-3 coronal brain sections per mouse were manually segmented in ImageJ, and the median immunofluorescence signal for each region of interest was measured. Prior to statistical analysis, values were normalized to the average value of all measurements across CAR T and mock treated mice within one experimental group within the same cage.

### Quantification of ICAM-1 and VCAM-1 expression in brain capillaries by immunofluorescence

Mice were euthanized on day 6 after CAR T cell infusion by CO2 inhalation and perfused transcardially with PBS followed by 4% paraformaldehyde and brains were dissected out. 40-50μm thick floating brain sections were incubated in blocking solution (10% donkey serum+0.1% Triton X-100 in PBS) for 1 h, then at 4°C overnight with primary antibodies directed against ICAM-1 (clone YN1/1.7.4, Biolegend, 1:100) or VCAM-1 (clone 429, Biolegend, 1:100) and laminin (rabbit polyclonal, Novus Biologicals cat# NB300-144, 1:50) in 1% donkey serum+0.1% Triton X-100 in PBS, washed in PBS, incubated with secondary antibodies for 2h at room temperature, counterstained with DAPI and mounted in FluoromountG (Southern Biotech). Stained sections were imaged with a confocal microscope at 20X and Z-projected as a maximum intensity projection. To measure expression of ICAM-1 and VCAM-1 in capillaries, regions of interest (ROIs) were defined by manually tracing each capillary segment in the laminin channel and measuring the brightness of the ROI in the ICAM-1 or VCAM-1 channel. Background brightness was measured by copying each ROI to an adjacent area without a vessel and again measuring the brightness of the ICAM-1 or VCAM-1 channel. The background signal was subtracted from each within-capillary measurement to obtain the final brightness per capillary segment. Two 466×466µm images of somatosensory cortex were analyzed per mouse. The average brightness per pixel across all capillaries was determined for each mouse, and normalized to the average brightness for all capillaries for all mice within the same experimental group.

### LFA-1 and VLA-4 affinity and expression in circulating leukocytes

Soluble fluorescent ICAM-1 and VCAM-1 multimers were produced by following a protocol adapted from Dimitrov et al (*54*). We incubated 200 µg/ml recombinant mouse ICAM-1-Fc chimera protein (Cat# 553006, Biolegend) or 200 µg/ml recombinant mouse VCAM-1-Fc chimera protein (Cat# 643-VM-050, R&D Systems) with polyclonal anti-human Fc-PE F(ab’)2 fragments (Cat# 109-116-170, Jackson Immuno Research) diluted 3.41x to final working volume with PBS. The multimer mixture was incubated at 4°C for 18 hours on a rocking shaker. As an example, in order to produce 100µl of ICAM-1 multimers, we mixed 20µg of ICAM-1-Fc with 29.3µl Fc-PE F(ab′)2 antibody (3.41x dilution) and diluted the mixture to 100µl final volume with PBS.

We obtained whole blood samples from mice across all treatment groups (saline control, cyclophosphamide only, mock transduced T cell treated, CAR T cell treated) via chin bleed on Days -1, 5, 12, and 19 post treatment. Blood samples were processed within 1 hour after collection. Spleens were dissected out and dissociated into a single cell suspension by mashing them through a 70μm filter. To measure the quantity and activation of integrins LFA-1 and VLA-4 on the surfaces of various leukocytes, fresh blood samples were diluted 1:1 with PBS and then transferred into a 96-well U-bottom plate at 20µl per well. After Fc blocking with anti-mouse CD16/32 (Cat# 101319, Biolegend) for 5 min, samples were stained with the following immunophenotyping antibodies: CD45.2 (Clone 104, Cat. 109832, Biolegend), CD3 (Clone 17A2, Cat. 100216, Biolegend), CD19 (Clone 6D5, Cat. 115530, Biolegend), CD11b (Clone M1/70, Cat. 101212, Biolegend), Ly6G (Clone 1A8, Cat. 127616, Biolegend), CD90.1 (Clone OX-7, Cat. 202503, Biolegend), CD8a (Clone 53-6.7, Cat. 100744, Biolegend), CD44 (Clone IM7, Cat. 103031, Biolegend), CD62L (Clone MEL-14, Cat. 104417, Biolegend) and Live/Dead Aqua (Cat. L34965, Thermo Fisher). CD11a (Clone M17/4, Cat. 101107) and CD49d (Clone PS/2, Cat. 1520-09L, Southern Biotech) were used to detect surface integrins LFA-1 and VLA-4, respectively, while ICAM-1 and VCAM-1 multimers were used to specifically bind to activated integrins. Samples were stained for 15 min then were fixed and lysed for erythrocytes with fix/lyse buffer (Cat. 00-5333-54, Thermo Fisher) for 10 min. Cells were washed with autoMACS buffer (Cat. 130-091-221, Miltenyi Biotec) twice before running flow cytometry.

### In vivo integrin blockade

Integrin αL function blocking antibody (Clone M17/2, Cat. BE0006, InVivoMAb), integrin α4 function blocking antibody (Clone PS/2, Cat. BE0071, InVivoMAb), or isotype control (clone LTF-2, Cat. BE0090, InVivoMAb) were retro-orbitally injected at a dose of 2mg/kg under isoflurane anesthesia on Days 1, 3, and 5 post treatment. Antibody doses were chosen to achieve a ∼25-50 μg/mL blood level to approximate the blood levels measured in human patients receiving natalizumab (*30*). On day 6, open field testing was conducted in a clean 20cm x 20cm x 20cm testing chamber. Video was acquired with a tracking camera above the chamber and analyzed with Noldus software. After a 5 minute acclimation period, 10 minutes of activity tracking were performed, and time spent in the center 50% of the arena and total distance traveled were measured. For statistical analysis, each mouse’s mean velocity during the 10 minute tracking period was converted into a percentage of the average of the velocity of all mice within the same cage who were part of the same experiment.

### 2D human brain microvascular endothelial cell ICAM-1 and VCAM-1 expression in response to cytokine stimulation

Primary human brain microvascular endothelial cells (HBMECs) were obtained from ScienCell Research Laboratories (cat# 1000) and cultured in flasks coated with 2% gelatin (Sigma). HBMECs from passages 3-5 were used for experiments. They were maintained in microvascular endothelial cell growth medium (cat# CC-3202, Lonza) that includes 5% fetal bovine serum (FBS), hydrocortisone, human fibroblast growth factor-beta (hFGF-β), vascular endothelial growth factor (VEGF), R3-insulin-like growth factor-1 (R3-IGF-1), ascorbic acid, human epidermal growth factor (hEGF), and gentamicin sulfate-amphotericin (GA-1000). Cells were incubated at 37°C with 5% CO2 and media replaced every 2 days.

Recombinant human tumor necrosis factor (TNF), interferon-γ (IFN-γ), interleukin-2 (IL-2), interleukin-4 (IL-4), interleukin-6 (IL-6), interleukin-8 (IL-8), interleukin-10 (IL-10), interleukin-15 (IL-15), granulocyte-macrophage colony-stimulating factor (GM-CSF), and C-X-C motif chemokine ligand 10 (CXCL-10) were obtained from Peprotech and diluted in complete microvascular endothelial cell growth medium (cat# CC-3202, Lonza) to reach final concentrations of 10 pg/ml, 100 pg/ml, and 1000 pg/ml. Recombinant human interleukin-1β (IL-1β, Peprotech) was diluted in complete growth medium to reach final concentrations of 0.5 pg/ml,1pg/ml, 2.5 pg/ml, 5 pg/ml, 10 pg/ml, and 100 pg/ml. HBMECs were cultured in 2% gelatin coated 24-well plates to 70-80% confluency and then treated with 500 µL of media containing one cytokine at various concentrations for 24 hours. Treatments were given to duplicate wells.

Cytokine-treated HBMECs were lifted from 24 well plates with TrypLE (ThermoFisher Scientific), and 50,000 cells per well were placed into 96 well plates. Cells were first stained with Live/Dead fixable far red dead cell stain (cat# L34973, ThermoFisher Scientific) at room temperature and then stained with anti-human VCAM-1 (=CD106, clone STA, cat# 305806, Biolegend) and anti-human ICAM-1 (=CD54, clone HA58, cat# 353108, Biolegend) antibodies for 15 minutes on ice in the dark. Flow cytometry analysis was performed on a Novocyte flow cytometer where 10,000 cells were acquired and single cells were identified using forward scatter height versus area. Median fluorescence intensity of ICAM-1 and VCAM-1 was used for analysis.

### Human CD19-CAR T cell generation

A clinically validated CD19-CAR (*33*) with a truncated EGFR transduction marker was used for all human in vitro experiments. Cryopreserved healthy donor PBMCs were thawed into XVIVO medium (Lonza) supplemented with IL-2 (4.6 ng/ml), IL-7 (5 ng/ml), IL-15 (0.5 ng/ml), and IL-21 (1 ng/ml; Miltenyi Biotec) at a density of 1 × 10^6^ cells/ml and activated with Human T-Activator CD3/CD28 Dynabeads (ThermoFisher). On day 1, cells were concentrated to a density of 4 × 10^6^/ml and transduced with lentiviral vectors encoding CAR constructs at an MOI of 2 with protamine sulfate (25 μg/ml). Twenty-four hours later, the cells were moved to GREX 24-well plates (Wilson Wolf) for further expansion at 1 × 10^6^/ml in XVIVO medium with cytokines until Dynabeads were removed by magnet on day 7. Cells were expanded until harvest on day 10-14.

### 3D human brain microvessel model fabrication

Primary HBMECs were purchased from Cell Systems (ACBRI 376), plated in a T-75 culture flask coated with Attachment Factor (4Z0-210, Cell Systems), and expanded in monolayer culture until passage 4-5 for seeding into microvessels. 3D microvessels were fabricated in 7.5 mg/mL type I collagen using soft lithography and collagen injection molding techniques as described previously(*55*). Briefly, after collagen injection and gelation, the top collagen pieces containing a micro-patterned, double-hexagonal branched network were assembled with flat bottom collagen pieces to form enclosed, perfusable 3D lumens within an acrylic housing system. Each microvessel was connected to inlet and outlet media reservoirs on the top housing jig. To uniformly endothelialize the microvessels, 10 µL of HBMEC suspension at 10×10^6^ cells/mL seeding density was perfused through each inlet and outlet reservoir under gravity-driven flow. Microvessels were cultured under gravity-driven perfusion of EGM-2MV media (Lonza) for 4-6 days with media replenishment every 12 hours to establish a confluent endothelial lumen. Starting on day 4-6, the media was supplemented with 100 pg/mL of each TNF and IFN-γ for 18 hours, followed by a 15-minute wash with cytokine-free media prior to the leukocyte perfusion studies. Control vessels were cultured in cytokine-free media throughout.

### Human CAR T cell perfusion studies in 3D brain microvessels

Cryopreserved human CD19-CAR T cells were thawed, rested for 2h, and resuspended at 1×10^6^ cells/mL in EGM-2MV media. Streptavidin plates (Thermo, cat # 15124) were coated with 5µg/mL biotinylated human CD19 (R&D, cat # AVI9269) in PBS overnight, and washed twice prior to plating CAR T cells in the wells for 18h of stimulation. Non-stimulated CAR T cells underwent the same procedure in uncoated wells. Following stimulation, 150 µL of CAR T cell suspension was perfused into cytokine-treated or control microvessels via the inlet reservoir followed by 50 µL of EGM-2MV media added to the outlet reservoir to establish 1 hour of gravity-driven flow. Unbound CAR T cells were removed by two 15-minute media washes. Microvessels were then fixed in 3.7% paraformaldehyde and washed with DPBS three times. Microvessels were incubated in blocking solution (2% bovine serum albumin and 0.1% Triton-X in DPBS) for 1 hour. Blocked microvessels were then incubated with conjugated antibodies against CD3 (Biolegend, clone HIT3a, 1:50), CD45 (BD Pharmingen, clone HI30, 1:50), and Hoechst 33342 (Thermo Fisher Scientific, 1:250) at 4°C overnight followed by three DBPS washes.

For adhesion blockade studies, CD19-stimulated CAR T cells were incubated with blocking antibodies against integrin αL (anti-LFA-1/CD11a, clone TS1/22, Genesee, cat # 55-8011U), integrin α4 (natalizumab, MedChem Express, cat # HY-108831A) alone or in combination, or an isotype control (clone S228P, Biocell, cat #CP147). Each antibody was added at 50 µg/mL in EGM-2MV media for 20 minutes prior to and during the 1 hour-perfusion in cytokine co-stimulated 3D microvessels. CAR T cell perfusion, media washes, microvessel fixation, blocking and immunostaining were performed as described above.

After staining, microvessels were imaged using a Nikon Ti2 microscope equipped with a Yokogawa W1 spinning disk confocal system. Z-stack tile scans were acquired using a 20x objective with a 2.5 µm z-step size to reconstruct the double-hexagonal branched network. For single-field imaging of the hexagonal branches, z-stacks were obtained using a 10x objective with a 5 µm z-step size. Bound human CD3+ CD45 + CAR T cells were quantified using maximum intensity projection images of the bottom luminal surface. Nuclei staining was used to verify mononuclear cell identity. “Multi-Point Tool” in ImageJ was used for cell counting.

### Human samples and analysis

Deidentified serum and plasma samples were obtained from patients with acute lymphoblastic leukemia who received CD19-directed CAR T cells as part of clinical trials at Seattle Children’s (NCT02028455, NCT03330691) (*33, 35*)or Fred Hutch Cancer Center (NCT01865617)(*32*). To measure soluble VCAM-1 and soluble ICAM-1, the VPLEX Vascular Injury Panel II (cat # K151198D-2, Meso Scale Diagnostics) was run and analyzed according to manufacturer’s instructions with a 1:10,000 sample dilution. Pooled healthy serum with spiked-in standard was used on all plates as the inter-plate control; all assays used in the analysis had interplate control coefficient of variation (CV) of 3-14%. Serum cytokines were measured using Milliplex (Millipore) or Meso Scale Discovery (Meso Scale Diagnostics) assays per the manufacturers’ instructions.

### Single-cell data analysis

Single cell gene expression data from patients treated with commercial CD19-directed CAR T cells (GEO accession GSE197268) (*36*) was queried for gene expression levels of ITGA4 (integrin α4) and ITGAL (integrin αL) in CAR T cells and native T cells. Patients in the dataset were included if they had a complete response to CAR T cells, and single cell sequencing data was available for either both the infusion product and sorted CAR T cells on day 7 post infusion (Patients 7, 8, 10, 11, 15, and 19 received axicabtagene ciloleucel; patients 22, 26, 27, 29, 30, and 32 received tisagenlecleucel), and/or paired PBMC samples from baseline and day 7 (Patients 6, 8, 10, 11, 12, 13, 15 received axicabtagene ciloleucel; patients 19, 21, 22, and 30 received tisagenlecleucel). Each sample was individually clustered in Seurat using a standard single cell workflow. Cells were filtered for less than 15% mitochondrial RNA transcripts and a minimum of 200 genes. Data was log normalized for each cell as ln(1+10^4*unique molecular identifier (UMI) count for the gene of interest/total UMI count for the cell). For every sample, all T cell clusters were included in pseudobulk gene expression analysis. For each gene of interest, the summed number of log normalized UMIs in the population of interest was divided by the number of cells in the population to obtain a mean expression level per cell.

### Study approval

All human studies were conducted with the approval of the Seattle Children’s and Fred Hutch Cancer Center Institutional Review Boards. The reanalysis of publicly available single cell RNA sequencing data (*36*) used only deidentified data and does not meet the definition of human subjects research. All mouse studies were approved by the Seattle Children’s Research Institute Animal Use and Care Committee.

### Statistical analysis

All analyses were performed blinded to the treatment group. Where appropriate, normality tests were performed prior to statistical tests. When normality was met, parametric analysis (t-test, one-way ANOVA with Holm–Sidak post-test to account for multiple comparisons, Pearson’s correlation) was performed and means were reported. Otherwise, nonparametric analyses (Mann–Whitney test, Spearman correlation) were performed and the median values were reported. The specific test used is indicated in the text and/or figure legends. Differences were considered statistically significant at the 95% confidence level. All statistical analyses were performed in GraphPad Prism.

## List of Supplementary Materials

Figs. S1 to S9

Table S1

Movies S1 to S3

## Acknowledgements

We thank Dr. Stoyan Dimitrov for technical advice on integrin activation measurement.

## Funding

National Institutes of Health K08NS118138 (JG)

National Institutes of Health R37CA275954 (JG)

National Institutes of Health R37CA26677 (HHG)

National Institutes of Health P30CA015704 (HHG)

American Cancer Society/St. Baldrick’s Foundation 933283 (HHG)

Norcliffe Foundation Center for Integrative Brain Research at Seattle Children’s

Research Institute (JG)

## Author contributions

Conceptualization: LP, YTT, RH, HKL, LDF, CJT, RAG, HHG, JG

Methodology: LP, YTT, RH, HKL, LDF, AYS, HHG, YZ, JG

Investigation: LP, YTT, RH, HKL, LDF, KB, MB, EMF, IHD, AYH, CEA, CJT, RAG, HHG, YZ, JG

Visualization: LP, YTT, HKL, JG

Funding acquisition: SEPS, CT, HHG, YZ, JG

Supervision: SEPS, AYS, CJT, AYH, HHG, YZ, JG

Writing – original draft: LP, JG

Writing – review & editing: LP, YTT, HKL, CJT, RAG, HHG, JG

## Data and materials availability

All original data will be made available by the authors upon reasonable request.

---

**Fig. S1.**
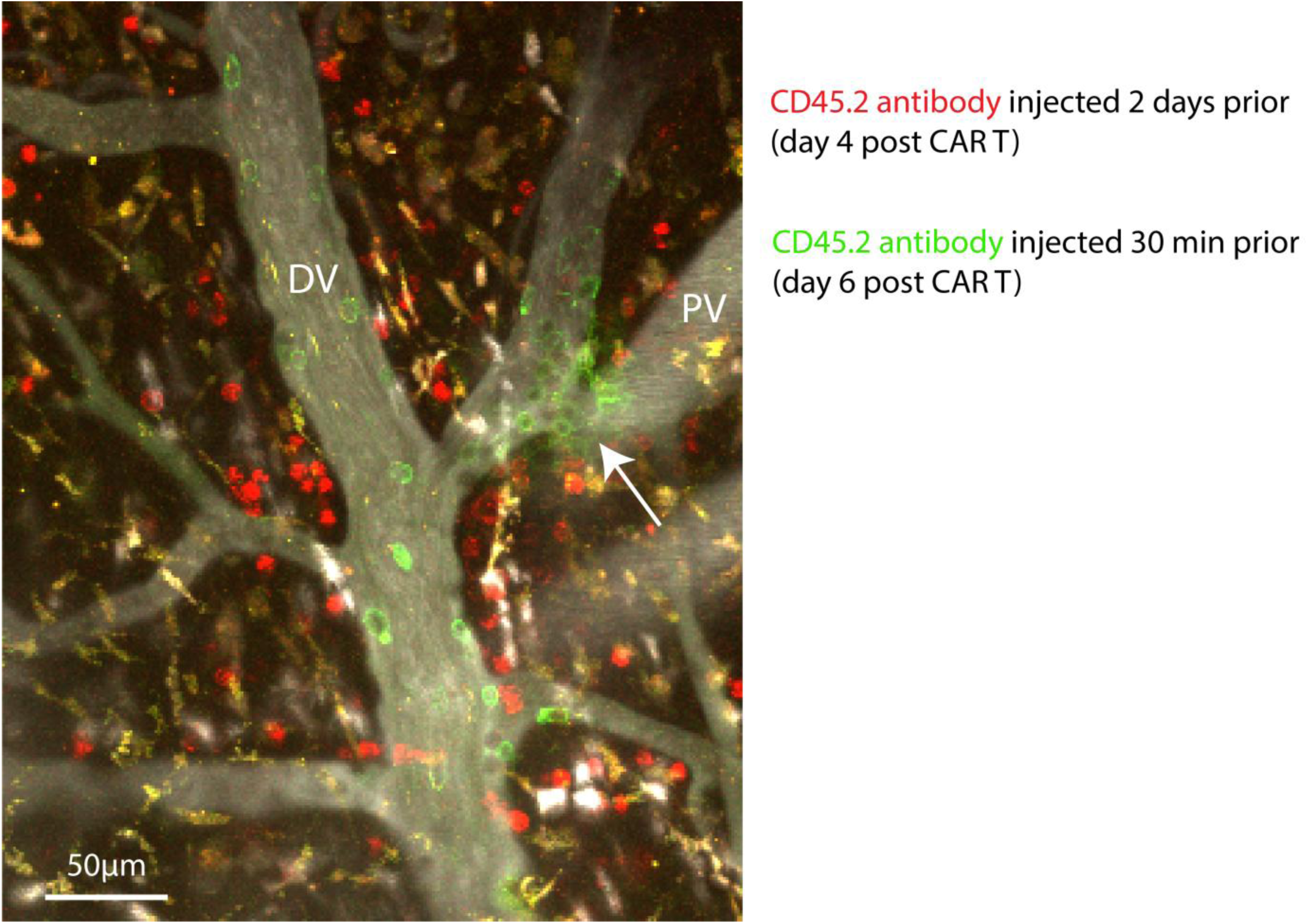
Active immune cell extravasation in the dura. In-vivo two-photon z-stack image of dural venule (DV, white intravascular dye) taken on day 6 after CAR T cell infusion. CD45.2 antibody conjugated to Alexa-568 (red) was given i.v. 2 days prior, and many labeled cells are seen spread throughout the dura. CD45.2 antibody conjugated to FITC (green) was injected 30 minutes prior to imaging, and labels primarily circulating leukocytes. The green intravascular cells are adherent to the wall of the venule. An active focus of extravasation is indicated by the arrow. All extravasated cells remained in the plane of the dura which is marked by autofluorescent (white/yellow) phagocytic cells that have taken up dyes from prior imaging. A pial venule (PV) is seen underlying the dura, it is not directly connected to the dural vasculature.

**Fig. S2.**
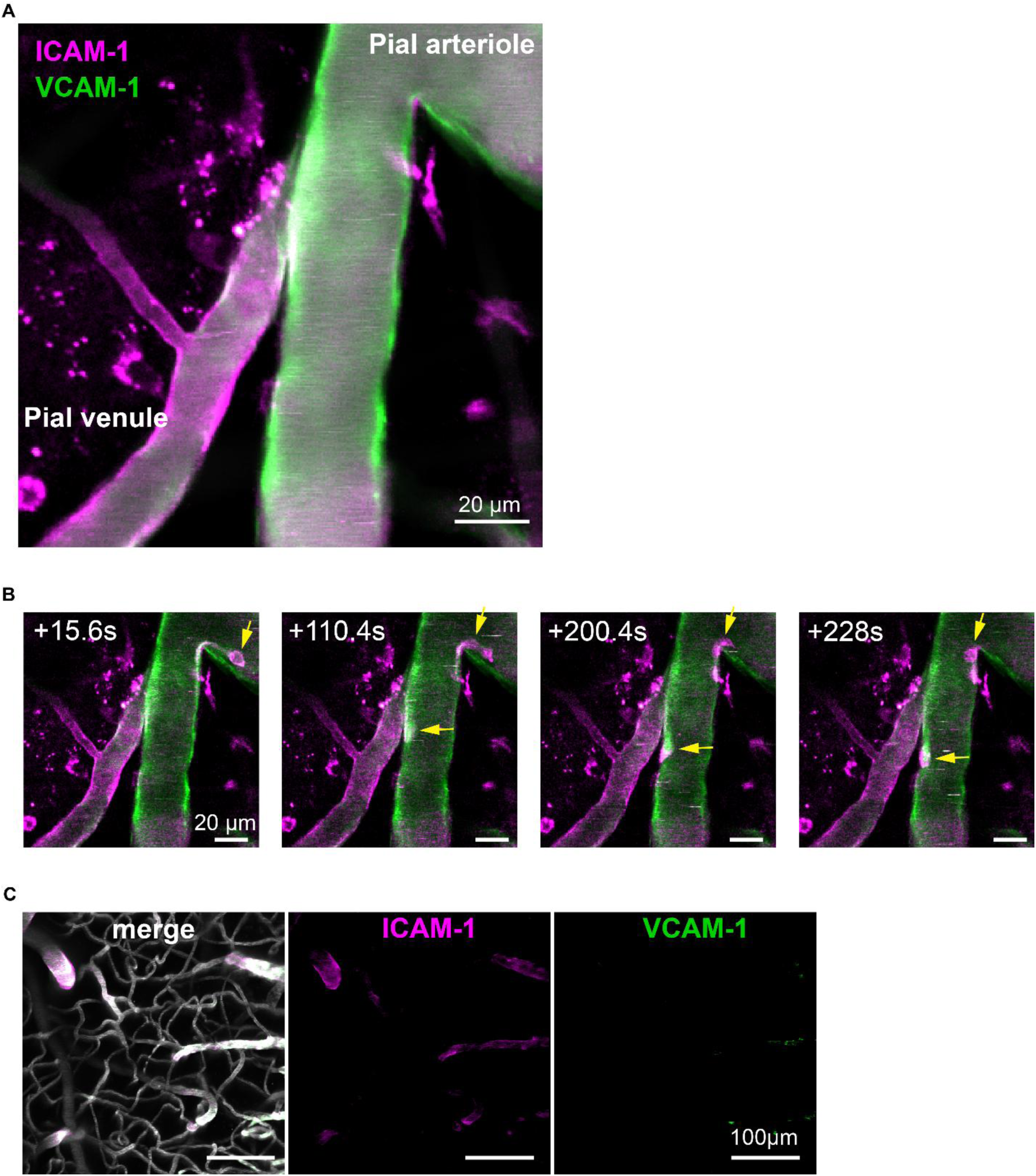
In vivo imaging of ICAM-1 and VCAM-1 expression. **(A)** strong ICAM-1 and VCAM-1 in pial vessels in a CAR T cell treated mouse. **(B)** Time lapse images of the same vessels as in (A), showing two leukocytes (arrows) crawling along the arteriolar wall. Also see Supplemental movie 2. **(C)** Low level ICAM-1 and very little VCAM-1 expression is seen in the pial vasculature in a control mouse receiving only cyclophosphamide preconditioning. There is very little expression in the capillarie. Images collected on day 6 after cyclophosphamide.

**Fig. S3.**
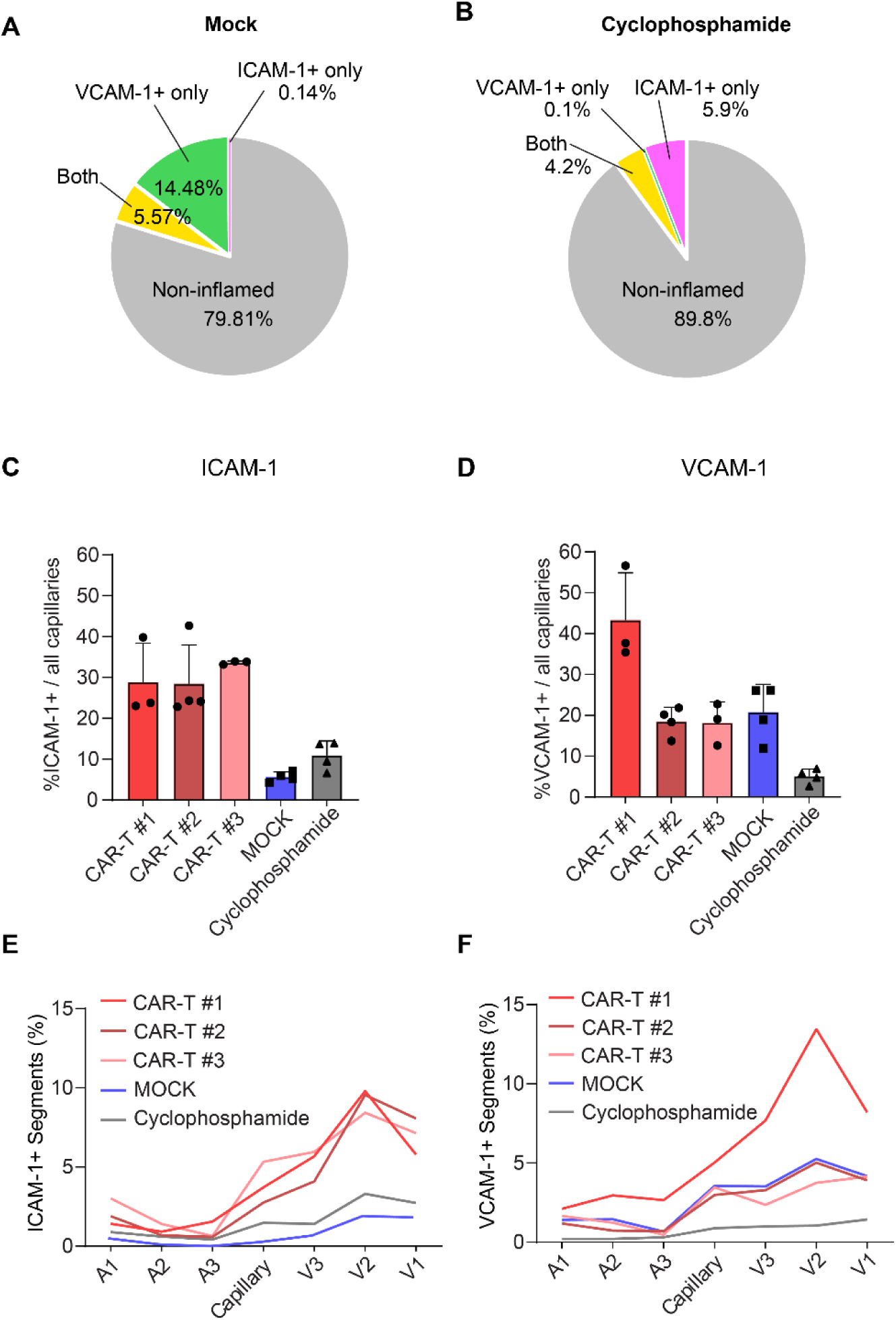
Capillary expression of ICAM-1 and VCAM-1 by in vivo two photon imaging. The distribution of in vivo antibody labeling is shown for mock **(A)** and cyclosphosphamide only **(B)** treated mice. N=1 mouse per condition, 4 imaging areas each, 718 capillary segments total (mock), 1000 capillary segments total (cyclophosphamide). The percentage of ICAM-1 **(C)** and VCAM-1 **(D)** positive capillaries is shown for each individual mouse that underwent imaging. Each spot is one imaging area, bars denote the mean and SEM. The distribution of ICAM-1 **(E)** and VCAM-1 **(F)** positive capillary segments is shown by branch order of brain parenchymal microvessels. The y-axis shows the percentage of all positive segments. Each line represents one mouse.

**Fig. S4.**
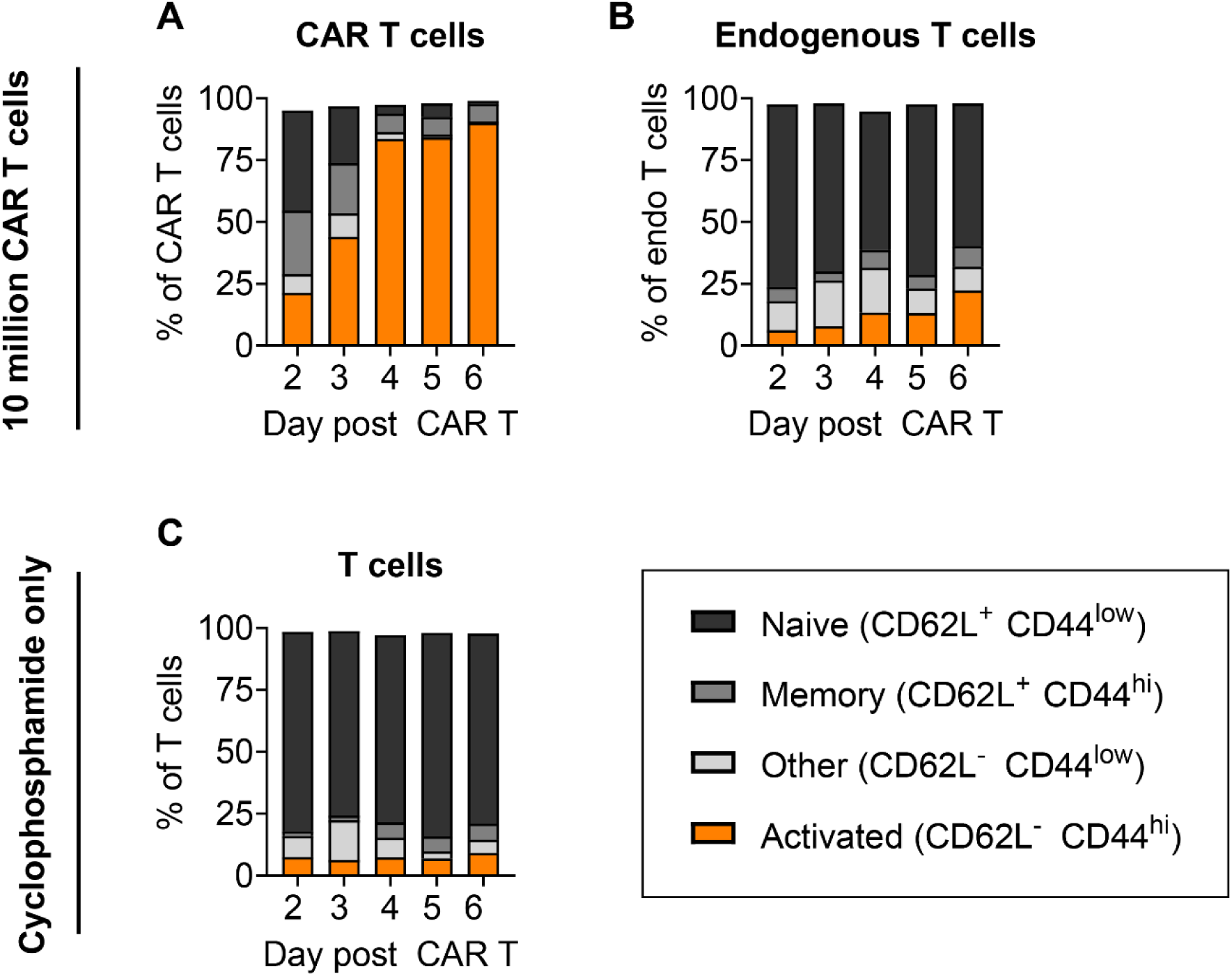
CAR T and endogenous T cell activation parallels kinetics of intravascular adhesion. Plots show T cell activation states defined by expression of CD62L and CD44 as measured by flow cytometry on dissociated splenocytes. **(A, B)**: mice received lymphodepletion on day -1 and CD19-CAR T cells on day 0; **(C)**: mice received cyclophosphamide lymphodepletion on day -1 only. Bars show the average percentage of each activation group (indicated in the legend) of all CAR T cells **(A)**, non-CAR endogenous T cells in mice treated with CAR T cells **(B)**, or all T cells in mice which did not receive CAR T cells **(C)**. 3 separate experiments, 3 mice total per time point. Totals may not add up to 100% due to unclassified T cells.

**Fig. S5.**
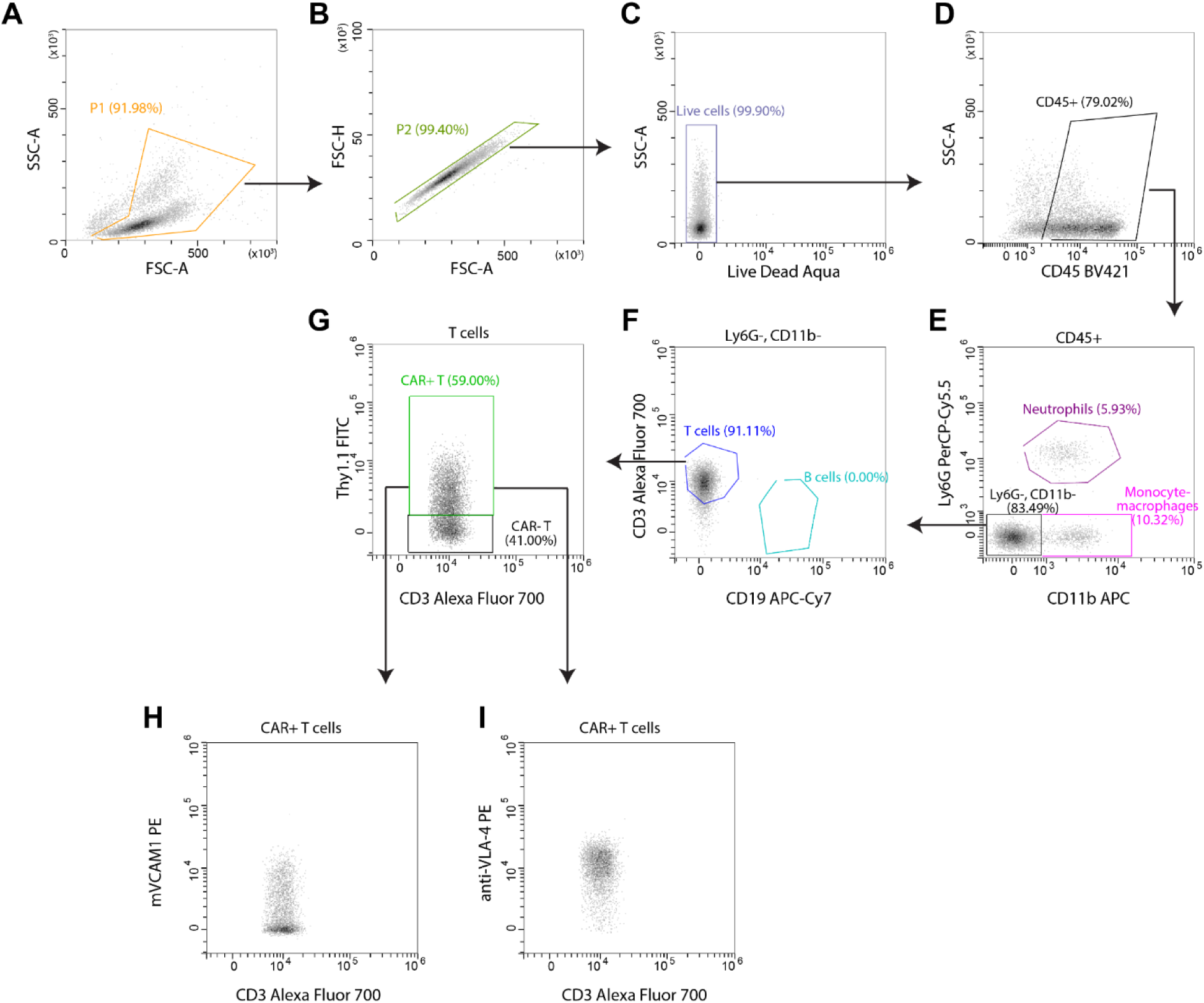
Gating strategy used for measuring median fluorescence intensity of VLA-4 affinity and surface VLA-4 within CAR+ T cell subset. Representative density plots from top left, following the arrows: Lymphocyte population (**A**) is gated for doublet exclusion (**B**), then further defined for live, CD45+ cells (**C,D**). After the exclusion of CD11b+ and Ly6G+ events (**E**), the CD3+ T cell population is obtained on CD3 vs CD19 (**F**). The resulting CD3+ population is further defined for CAR+ and CAR- T cells. CAR+ gate can be used to plot against mVCAM1 or VLA-4 to measure the mean fluorescence intensity of VLA4 affinity or expression, respectively.

**Fig. S6.**
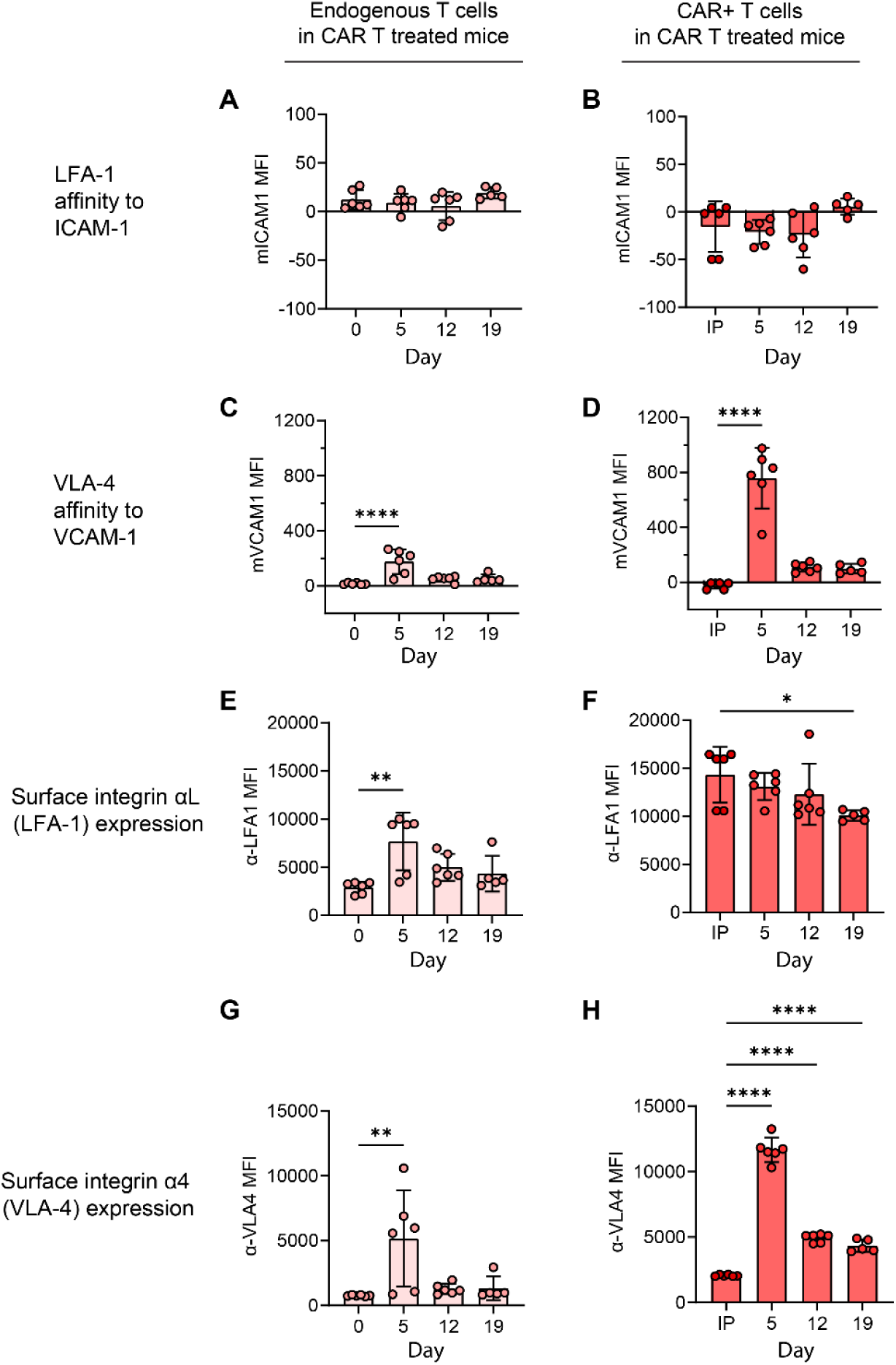
Kinetics of VLA-4 and LFA-1 affinity and expression in CAR T cell treated mice. Data for endogenous, untransduced circulating T cells in shown in the left column, and data for circulating CAR T cells (or infusion product) is shown in the right column. The bar graphs show flow cytometry measurements of T cell affinity to ICAM-1 (**A,B**) or VCAM-1 (**C,D**) multimers, as well as cell surface expression of integrins αL (**E,F**) and α4 (**G,H**). The x-axis shows days relative to CAR T cell infusion (IP, cryopreserved infusion product). The y-axis shows mean fluorescence intensity (arbitrary units). Each data point indicates one mouse, the box plots show the mean and the whiskers show the standard deviation. N=3 independent experiments, for a total of 6 mice. One-way ANOVA with Dunnett’s multiple comparisons test. Only comparisons with Day 0 (left column) or IP (right column) were performed. For clarity, only statistics with P<0.05 are shown. *P<0.05, **P<0.01, ***P<0.001, ****P<0.0001.

**Fig. S7.**
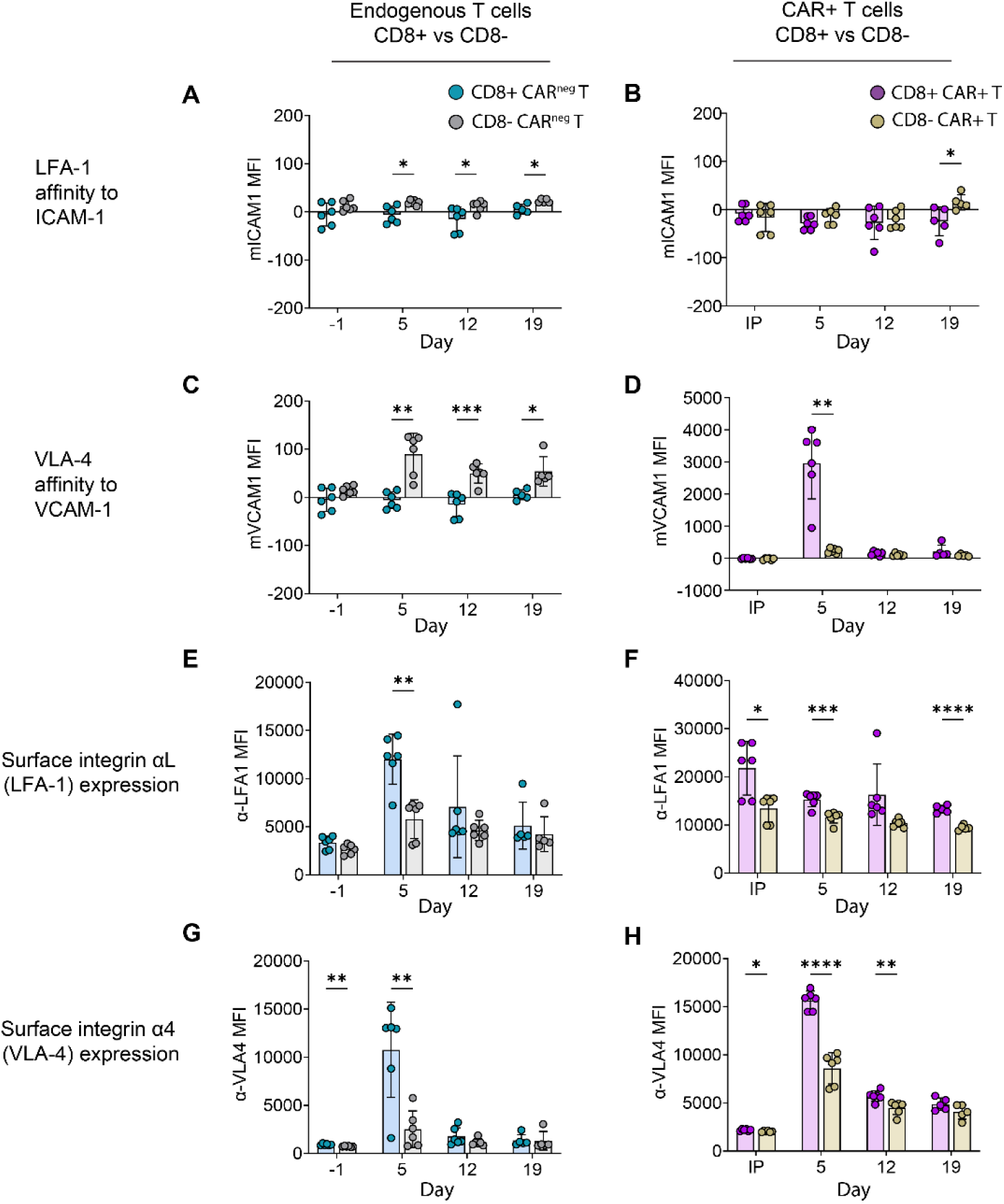
Kinetics of VLA-4 and LFA-1 affinity and expression differ between T cell subsets. Data for endogenous untransduced circulating T cells is shown in the left column (**A,C,E,G**), and data for circulating CAR T cells (or infusion product) is shown in the right column (**B,D,F,H**). The bar graphs compare flow cytometry measurements between CD8+ and CD8- T cells. In endogenous T cells, the CD8- (=CD4+) subset had higher affinity to ICAM-1 (A) and VCAM-1 (C) compared to CD8+ cells. Neither subset of CAR T cells had a change in ICAM-1 affinity (B). Only CD8+ CAR T cells strongly increased their affinity to VCAM-1 (D). Cell surface expression of integrins αL (E,F) and α4 (G,H) was generally highest in CD8+ T cells, but CD8-T cells showed a similar temporal pattern. The x-axis shows days relative to CAR T cell infusion (IP, cryopreserved infusion product). The y-axis shows mean fluorescence intensity (arbitrary units). Each data point indicates one mouse, the box plots show the mean and the whiskers show the standard deviation. N=3 independent experiments, for a total of 6 mice. Multiple unpaired t test with Welch correction. For clarity, only statistics with P<0.05 are shown. *P<0.05, **P<0.01, ***P<0.001, ****P<0.0001.

**Fig. S8.**
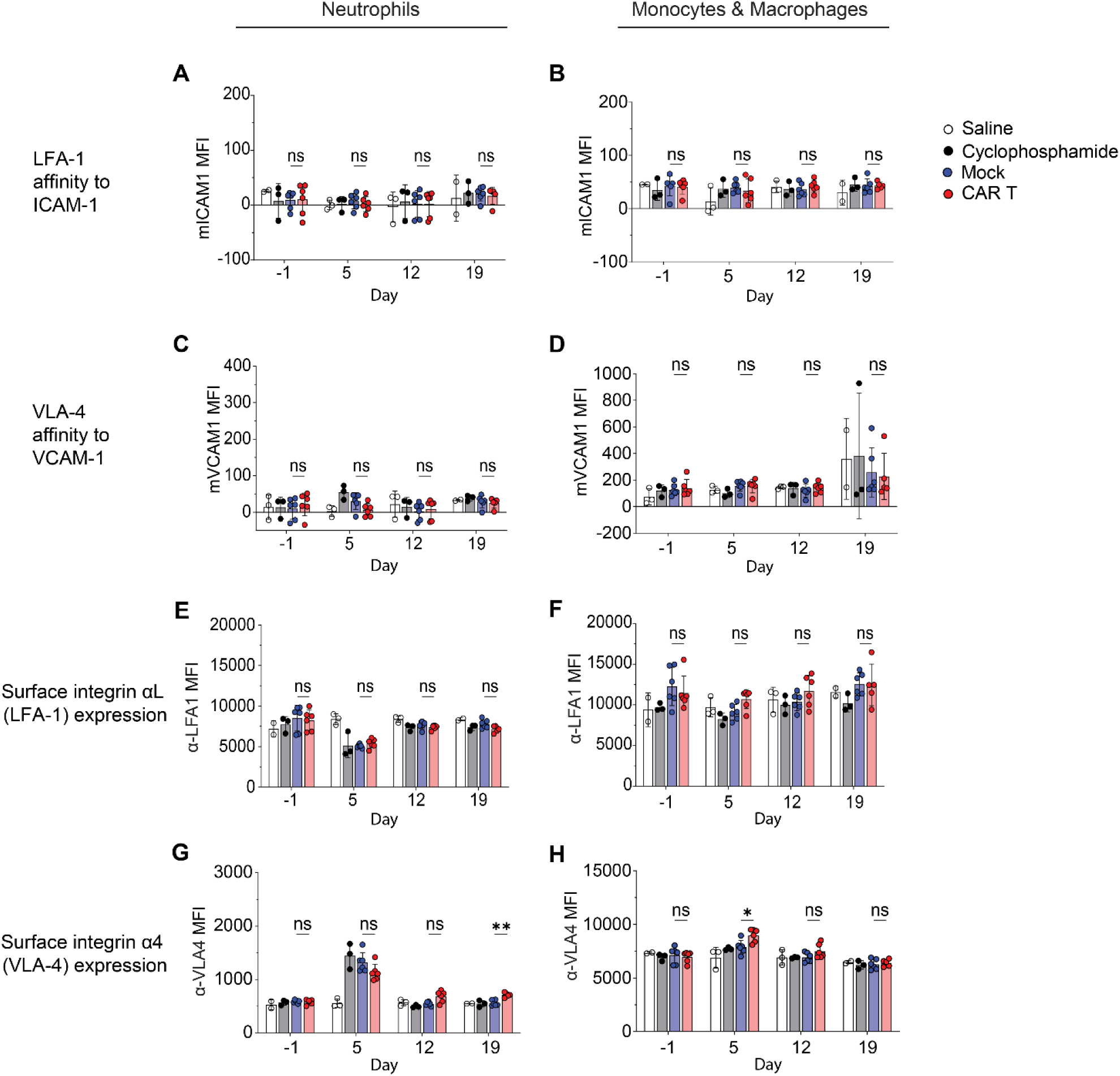
Circulating neutrophils and monocytes/macrophages have minimal modulation of VLA-4 and LFA-1 affinity and expression after CAR T cell treatment. Scatter plots compare flow cytometry measurements of neutrophil (left column) or monocyte/macrophage (right column) affinity to ICAM-1 (**A,B**) or VCAM-1 (**C,D**) multimers, as well as cell surface expression of integrins αL (**E,F**) and α4 (**G,H**). The y-axis indicates the mean fluorescence intensity (MFI, arbitrary units) for each analyte, and the x-axis indicates time relative to CAR T cell (or control) infusion. Each data point indicates one mouse, the box plots show the mean and the whiskers show the standard deviation. N=3 independent experiments, for a total of 3-6 mice per group. All comparisons are by unpaired 2-sided t test, *P<0.05, **P<0.01, ***P<0.001, ****P<0.0001, ns P>0.05. For clarity, only statistics comparing mock to CAR T treated mice are shown.

**Fig. S9.**
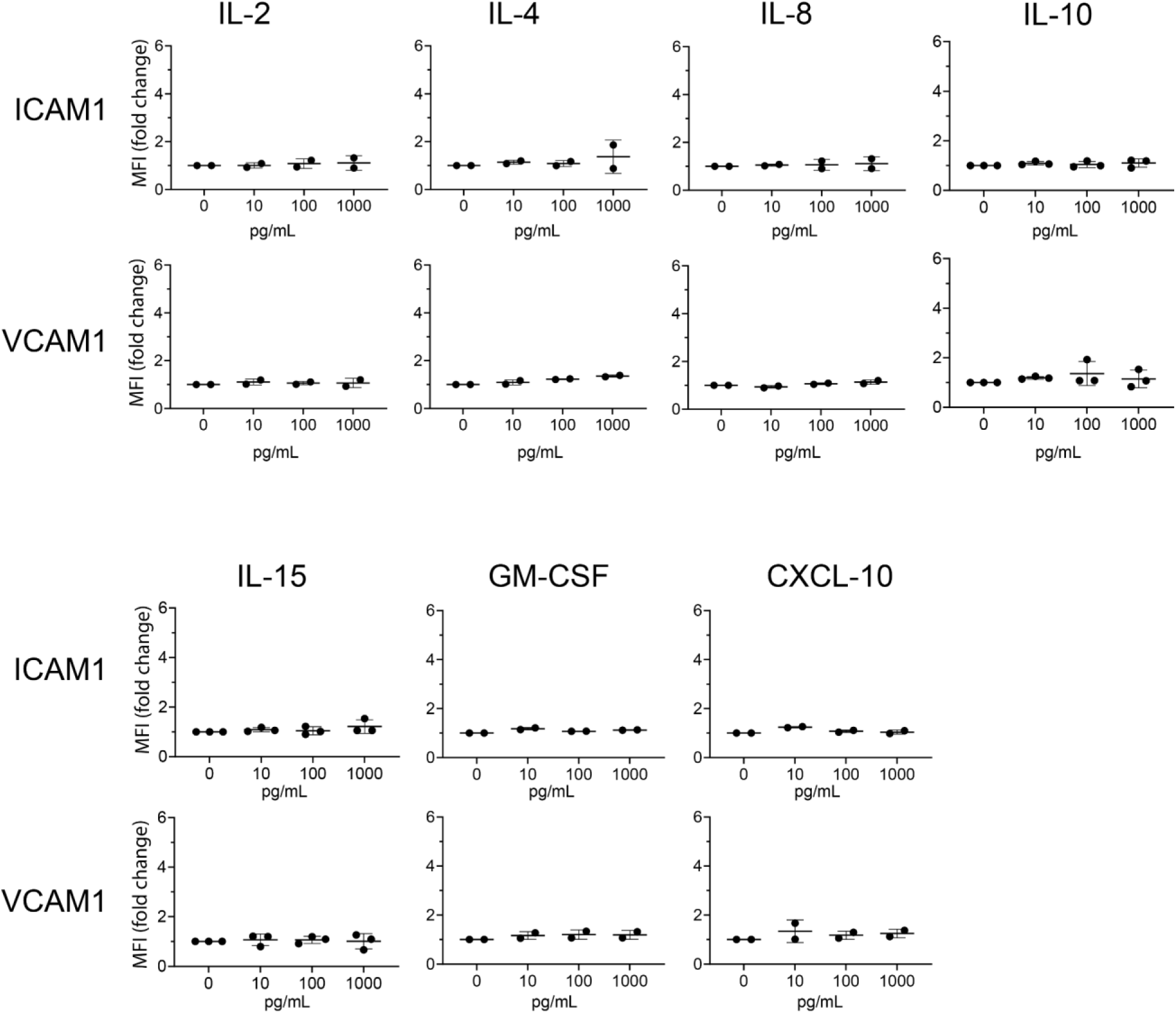
Most ICANS-associated cytokines do not increase brain microendothelial expression of ICAM-1 or VCAM-1. Scatter plots show surface expression of ICAM-1 and VCAM-1 in hBMECs after treatment with cytokines. The y-axes indicate the fold change in mean fluorescence intensity as measured by flow cytometry, comparing each data point to the 0 pg/mL vehicle-only control within the same experiment. The x-axis indicates the concentration of each cytokine that was added to the culture media for 24h. N=3 independent experiments, one-way ANOVA with Dunnett’s multiple comparisons test, only comparisons with the 0 pg/mL vehicle-only control were performed. All comparisons were not significant with P>0.05.

**Table S1.** Soluble ICAM-1 and VCAM-1 levels in patient plasma. Mean (top table) or median (bottom table) values are shown. “Day” indicates day after CAR T cell infusion. All statistics compare patients with ICANS (grade 1-5) to patients with no ICANS (grade 0) on the same day post CAR T within the same clinical trial. No adjustment was made for multiple comparisons.

|  | ICANS | N= | Statistic | soluble ICAM-1 |  |  |  |  | soluble VCAM-1 |  |  |  |
| --- | --- | --- | --- | --- | --- | --- | --- | --- | --- | --- | --- | --- |
|  |  |  |  | Day 1<br>(ng/mL) | Day 7<br>(ng/mL) | Day 7/<br>Day 1 | Day 7-Day 1 |  | Day 1<br>(ng/mL) | Day 7<br>(ng/mL) | Day 7/<br>Day 1 | Day 7-Day 1 |
| Adult CD19-CAR | yes | 24 | mean | 547.0 | 950.4 | 1.698 | 403.3 |  | 959.5 | 1271.3 | 1.307 | 311.8 |
|  | no | 24 | mean | 693.1 | 827.7 | 1.058 | 29.3 |  | 877.8 | 1065.2 | 1.152 | 147.1 |
|  |  |  | *P (unpaired t test)* | *0.0884* | *0.5882* | *0.0462* | *0.0403* |  | *0.5846* | *0.3137* | *0.0435* | *0.0542* |
| Pediatric<br>CD19-CAR | yes | 18 | mean | 1024.5 | 1373.0 | 1.322 | 348.4 |  | 953.4 | 1153.7 | 1.272 | 200.4 |
|  | no | 19 | mean | 977.7 | 973.1 | 1.038 | -4.6 |  | 1086.1 | 1039.3 | 1.055 | -46.8 |
|  |  |  | *P (unpaired t test)* | *0.7717* | *0.1999* | *0.1085* | *0.1633* |  | *0.4342* | *0.4239* | *0.1427* | *0.0492* |
| Pediatric<br>CD19/CD22-CAR | yes | 7 | mean | 605.3 | 1071.1 | 1.735 | 465.8 |  | 688.6 | 1264.5 | 1.791 | 575.9 |
|  | no | 17 | mean | 707.1 | 909.4 | 1.279 | 202.3 |  | 828.6 | 1120.0 | 1.325 | 291.3 |
|  |  |  | *P (unpaired t test)* | *0.3316* | *0.5131* | *0.1112* | *0.206* |  | *0.3627* | *0.6434* | *0.0371* | *0.1569* |
|  |  |  |  | Day 1<br>(ng/mL) | Day 7<br>(ng/mL) | Day 7/<br>Day 1 | Day 7-Day 1 |  | Day 1<br>(ng/mL) | Day 7<br>(ng/mL) | Day 7/<br>Day 1 | Day 7-Day 1 |
| Adult CD19-CAR | yes | 24 | median | 508.7 | 641.6 | 1.270 | 132.8 |  | 779.5 | 995.9 | 1.223 | 196.3 |
|  | no | 24 | median | 555.9 | 626.6 | 1.107 | 48.7 |  | 823.3 | 895.8 | 1.155 | 102.6 |
|  |  |  | *P (Mann-Whitney)* | *0.2465* | *0.4811* | *0.0025* | *0.0045* |  | *0.9106* | *0.5585* | *0.0930* | *0.0814* |
| Pediatric<br>CD19-CAR | yes | 18 | median | 692.0 | 978.1 | 1.133 | 111.7 |  | 950.1 | 1087.9 | 1.084 | 86.3 |
|  | no | 19 | median | 918.5 | 865.5 | 0.978 | -20.0 |  | 797.6 | 1080.0 | 1.009 | 6.6 |
|  |  |  | *P (Mann-Whitney)* | *0.9163* | *0.2844* | *0.0656* | *0.0492* |  | *0.8689* | *0.4079* | *0.1259* | *0.1113* |
| Pediatric<br>CD19/CD22-CAR | yes | 8 | median | 610.5 | 866.2 | 1.455 | 221.9 |  | 669.3 | 1122.6 | 1.603 | 395.0 |
|  | no | 16 | median | 584.8 | 777.7 | 1.231 | 132.5 |  | 691.0 | 934.4 | 1.269 | 185.5 |
|  |  |  | *P (Mann-Whitney)* | *0.9284* | *0.8340* | *0.4523* | *0.4523* |  | *0.5283* | *0.3196* | *0.1056* | *0.1924* |

